# Cholinergic Neuronal Activity Promotes Diffuse Midline Glioma Growth through Muscarinic Signaling

**DOI:** 10.1101/2024.09.21.614235

**Authors:** Richard Drexler, Antonia Drinnenberg, Avishai Gavish, Belgin Yalcin, Kiarash Shamardani, Abigail Rogers, Rebecca Mancusi, Kathryn R. Taylor, Yoon Seok Kim, Pamelyn J. Woo, Alexandre Ravel, Eva Tatlock, Charu Ramakrishnan, Alberto E. Ayala-Sarmiento, David Rincon Fernandez Pacheco, La’Akea Siverts, Tanya L. Daigle, Bosiljka Tasic, Hongkui Zeng, Joshua J. Breunig, Karl Deisseroth, Michelle Monje

## Abstract

Neuronal activity promotes the proliferation of healthy oligodendrocyte precursor cells (OPC) and their malignant counterparts, gliomas. Many gliomas arise from and closely resemble oligodendroglial lineage precursors, including diffuse midline glioma (DMG), a cancer affecting midline structures such as the thalamus, brainstem and spinal cord. In DMG, glutamatergic and GABAergic neuronal activity promotes progression through both paracrine signaling and through bona-fide neuron-to-glioma synapses. However, the putative roles of other neuronal subpopulations - especially neuromodulatory neurons located in the brainstem that project to long-range target sites in midline anatomical locations where DMGs arise - remain largely unexplored. Here, we demonstrate that the activity of cholinergic midbrain neurons modulates both healthy OPC and malignant DMG proliferation in a circuit-specific manner at sites of long-range cholinergic projections. Optogenetic stimulation of the cholinergic pedunculopontine nucleus (PPN) promotes glioma growth in pons, while stimulation of the laterodorsal tegmentum nucleus (LDT) facilitates proliferation in thalamus, consistent with the predominant projection patterns of each cholinergic midbrain nucleus. Reciprocal signaling was evident, as increased activity of cholinergic neurons in the PPN and LDT was observed in pontine DMG-bearing mice. In co-culture, hiPSC-derived cholinergic neurons form neuron-to-glioma networks with DMG cells and robustly promote proliferation. Single-cell RNA sequencing analyses revealed prominent expression of the muscarinic receptor genes *CHRM1* and *CHRM3* in primary patient DMG samples, particularly enriched in the OPC-like tumor subpopulation. Acetylcholine, the neurotransmitter cholinergic neurons release, exerts a direct effect on DMG tumor cells, promoting increased proliferation and invasion through muscarinic receptors. Pharmacological blockade of M1 and M3 acetylcholine receptors abolished the activity-regulated increase in DMG proliferation in cholinergic neuron-glioma co-culture and *in vivo*. Taken together, these findings demonstrate that midbrain cholinergic neuron long-range projections to midline structures promote activity-dependent DMG growth through M1 and M3 cholinergic receptors, mirroring a parallel proliferative effect on healthy OPCs.

**HIGHLIGHTS:**

1. Activity of midbrain cholinergic neuron long-range projections increases proliferation of both healthy oligodendrocyte precursor cells (OPC) and malignant diffuse midline glioma (DMG) cells.
2. Optogenetic stimulation of cholinergic midbrain nuclei promotes growth in thalamic and pontine glioma in a circuit-dependent manner.
3. Reciprocally, DMG cells increase cholinergic neuronal activity in cholinergic midbrain nuclei.
4. The muscarinic receptors CHRM1 and CHRM3 are identified as therapeutic targets in DMG.

## INTRODUCTION

The activity of the nervous system has emerged as a crucial driver of cancers (for reviews, please see ^1–3^), and powerfully drives the progression^4–8^ and initiation^9,10^ of primary brain cancers called gliomas. High-grade gliomas, including glioblastoma and diffuse midline glioma (DMG) are the most common malignant brain tumor types and leading causes of brain-tumor-related death in adults and children^11^, respectively. DMGs, which are driven by oncogenic mutations in genes encoding histone H3 (H3K27M), originate most commonly in the pons of the brainstem and are also referred to as diffuse intrinsic pontine glioma (DIPG). H3K27M-altered DMGs also occur in the thalamus and spinal cord^12^. Regardless of their specific location in these midline structures, DMGs arise from and closely resemble oligodendrocyte precursor cells (OPCs)^12–17^, and numerous mechanistic parallels exist between the regulation of oligodendroglial lineage precursors and their malignant counterparts (for review, please see^18^). OPCs communicate extensively with neurons, both through neuron-to-OPC synapses^19,20^ and via paracrine factors such as brain-derived neurotrophic factor (BDNF)^21^. Within these neuron-to-OPC networks, neuronal activity promotes the proliferation of normal OPCs, oligodendrogenesis, and adaptive myelin changes that contribute to a range of functions including memory and learning in the healthy brain^21–26^ (for reviews on myelin plasticity please see^18,27^). In a similar pattern, neuronal activity also drives glioma proliferation^4–9,28^ and invasion^29,30^. The mechanisms mediating these growth-promoting effects of neurons on glioma cells include activity-regulated paracrine factor signaling^4,5,8^ and direct neuron-to-glioma synaptic communication^6–8^. Glutamatergic, AMPA receptor-mediated neuron-to-glioma synapses are found in both glioblastoma and DMGs^6–8,29^, while GABAergic neuron-to-glioma synapses are found selectively in DMGs^28^.

To date, research efforts have chiefly focused on the influence of glutamatergic and GABAergic neurons on glioma pathophysiology, while the effects of neuromodulatory neuron types - cholinergic, serotonergic, adrenergic and dopaminergic neurons - remain largely unexplored apart from the therapeutic potential of targeting dopamine receptor subtypes in glioblastoma^31^. Neural circuits are particularly salient to DMGs, given the brainstem location and robust axonal projections to midline structures of most neuromodulatory nuclei in the central nervous system. In particular, cholinergic midbrain nuclei such as the laterodorsal tegmentum nucleus (LDT) and pedunculopontine nucleus (PPN) project to midline structures where DMGs arise, highlighting the potential for these cholinergic neurons to modulate glioma pathophysiology. Here, we test this hypothesis and discover that cholinergic neuronal activity drives the proliferation of both healthy OPCs and DMG cells in a circuit-specific manner.

## RESULTS

### Cholinergic neuronal activity increases OPC proliferation

As neuronal activity influences both OPCs^23^ and DMGs^4–6,8^, and given the resemblance of DMG cells to OPCs^13,14,32^, we first explored the question of whether cholinergic neuronal activity modulates healthy OPC proliferation. We leveraged *in vivo* optogenetic stimulation to specifically target cholinergic midbrain neurons in either the pedunculopontine nucleus (PPN) or the laterodorsal tegmentum nucleus (LDT) (**Fig. 1A**), which respectively project to the pons and thalamus. To stimulate cholinergic neurons, we used ChAT-IRES-Cre mice cross-bred with Ai230 mice, a Cre-dependent reporter expressing the opsin ChRmine (ChAT-IRES-Cre^+/wt^ x Ai230^flx/wt^), thereby enabling stimulation of cholinergic neuronal activity using a potent opsin^33,34^ specifically expressed in cholinergic neurons (**Supplementary Fig. 1A**). We placed an optical ferrule within either the LDT (**Supplementary Fig. 1B**) or PPN (**Supplementary Fig. 1C**) to selectively stimulate cholinergic neurons confined to a single cholinergic nucleus. Light stimulation of the PPN or LDT at 20 Hz increased cFos expression in cholinergic neurons after 90 minutes, demonstrating successful optogenetic stimulation (**Supplementary Fig. 1D-H**). As cholinergic projections originating from the PPN and LDT target distinct brain regions, we performed optogenetic stimulation of each nucleus and evaluated OPC proliferation in response to cholinergic neuronal activity by administering the thymidine analogue EdU at the time of each stimulation session. As expected, predominant projections from LDT to thalamus (LDT->thalamus) and from PPN to pons (PPN->pons) were visualized utilizing Cre-dependent AAV tracing of cholinergic neurons (Supplementary Fig. 1F-G). In line with the anatomy of these projections, stimulation of cholinergic neurons in the LDT increased OPC proliferation in the thalamus (**Fig. 1B**), whereas PPN stimulation increased OPC proliferation in the pons (**Fig. 1C**), as indicated by quantification of EdU^+^/Pdgfra^+^ cells (**Fig. 1D**). Taken together, these results illustrate circuit-specific OPC proliferation consistent with target sites of midbrain cholinergic axonal projections.

**Figure 1.**
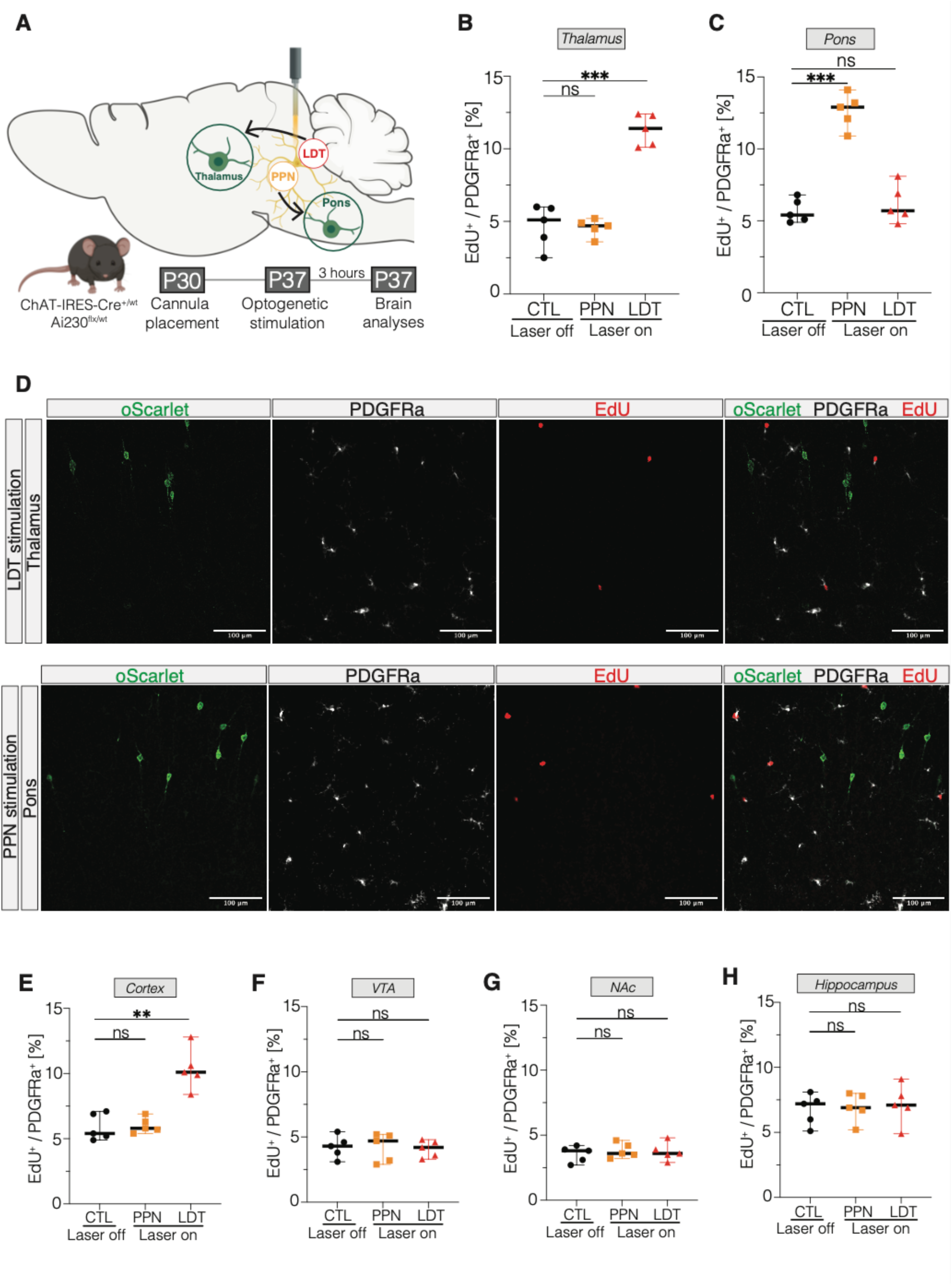
Cholinergic neuronal activity-regulated modulation of oligodendrocyte precursor cells. **(A)** Schematic of experimental paradigm for Optogenetic stimulation of cholinergic neurons in either laterodorsal tegmentum nucleus (LDT) or pedunculopontine nucleus (PPN). 5-week-old ChAT-IRES-Cre^+/wt^ x Ai230^flx/wt^ mice (P35-38) were stimulated for 30 minutes followed by perfusion after 3 hours. **(B)** Optogenetic stimulation of cholinergic neurons in LDT increases OPC proliferation (EdU^+^/Pdgfra^+^) in thalamus which is not seen after PPN stimulation (CTL (not stimulated), PPN stimulated, and LDT stimulated, n=5 mice/group). Unpaired two-tailed t-test; ***p < 0.001, ns: non-significant. Data shown as mean, error bars indicate range. **(C)** Optogenetic stimulation of cholinergic neurons in PPN increases OPC proliferation (EdU^+^/Pdgfra^+^) in pons (CTL, PPN, and LDT, n=5 mice). Unpaired two-tailed t-test; ***p < 0.001, ns: non-significant. Data shown as mean, error bars indicate range. **(D)** Confocal micrographs show thalamic OPC response after optogenetic stimulation of LDT (top image) and in pons after optogenetic stimulation of PPN (bottom image). ChRmine-oScarlet: green; Pdgfra: white; EdU: red, scale bars = 100um. **(E) – (H)** OPC response in **(E)** prefrontal cortex, **(F)** ventral tegmental area, **(G)** nucleus accumbens, and **(H)** hippocampus after optogenetic stimulation of cholinergic neurons (CTL, PPN, and LDT, n=5 mice/group). Unpaired two-tailed t-test; **p < 0.01, ns: non-significant. Data shown as mean, error bars indicate range.

Cholinergic axons from LDT and PPN also project to a variety of neuroanatomical sites. We found that LDT stimulation resulted in notably increased OPC proliferation in the cortex (**Fig. 1E**), with no changes in OPC proliferation observed in the ventral tegmental area (**Fig. 1F**), nucleus accumbens (**Fig. 1G**), or hippocampus (**Fig. 1H**). Despite the known functional and topographical organization of cholinergic projections to these regions, these data demonstrate a brain region-specific response of OPCs to cholinergic neuronal activity, consistent with the heterogeneous and region-specific OPC responses previously demonstrated for dopaminergic neurons^35^ in the VTA and glutamatergic cortical projection neurons^23^ in frontal cortex.

### Cholinergic neuronal activity modulates glioma growth

Given the similarities between OPCs and DMG, and the observed cholinergic circuit-specific thalamic and pontine OPC responses to cholinergic neuronal activity, we next asked whether DMG cells are equally responsive to long-range cholinergic neuronal activity in these midline structures. We addressed this by combining the previously mentioned genetically engineered optogenetic mouse model (ChAT-IRES-Cre^+/wt^ x Ai230^flx/wt^) with an *in utero* electroporation-induced genetic mouse model of H3K27M-DMG that leverages dual recombinase-mediated cassette exchange to express mutations in *H3f3a*, *Tp53* and *Pdgfra* in neural precursor cells (MADR)^36^. This model faithfully recapitulates an H3.3-K27M DMG^36^ and allows allografting into immunocompetent mice. The OPC-like characteristics of DMGs previously described in the MADR model^36^ were re-confirmed by *Pdgfra*-positivity in GFP^+^-tumour cells (**Supplementary Fig. 1I**).

Allografting into either the thalamus or pons was performed in 4-week-old mice (postnatal day (P) 28-30), with optical ferrule placement in the PPN or LDT three weeks later (P51). Optogenetic stimulation of cholinergic neurons was conducted after 4 weeks of tumour growth (P58), with administration of EdU at the time of the stimulation session. Mice were perfused 24-hours after optogenetic stimulation (**Fig. 2A**). As observed for OPCs, stimulation of cholinergic neurons in the LDT increased tumour cell proliferation in thalamic allografts more than 2-fold (**Fig. 2B,D**); no discernible effect was observed in the pons after LDT stimulation (**Fig. 2C**), consistent with the anatomy of cholinergic axon projections. Concordantly, stimulation of PPN cholinergic neurons increased DMG tumour cell proliferation in pontine allografts (**Fig. 2C**), but not in thalamic allografts (**Fig. 2B-C**).

**Figure 2.**
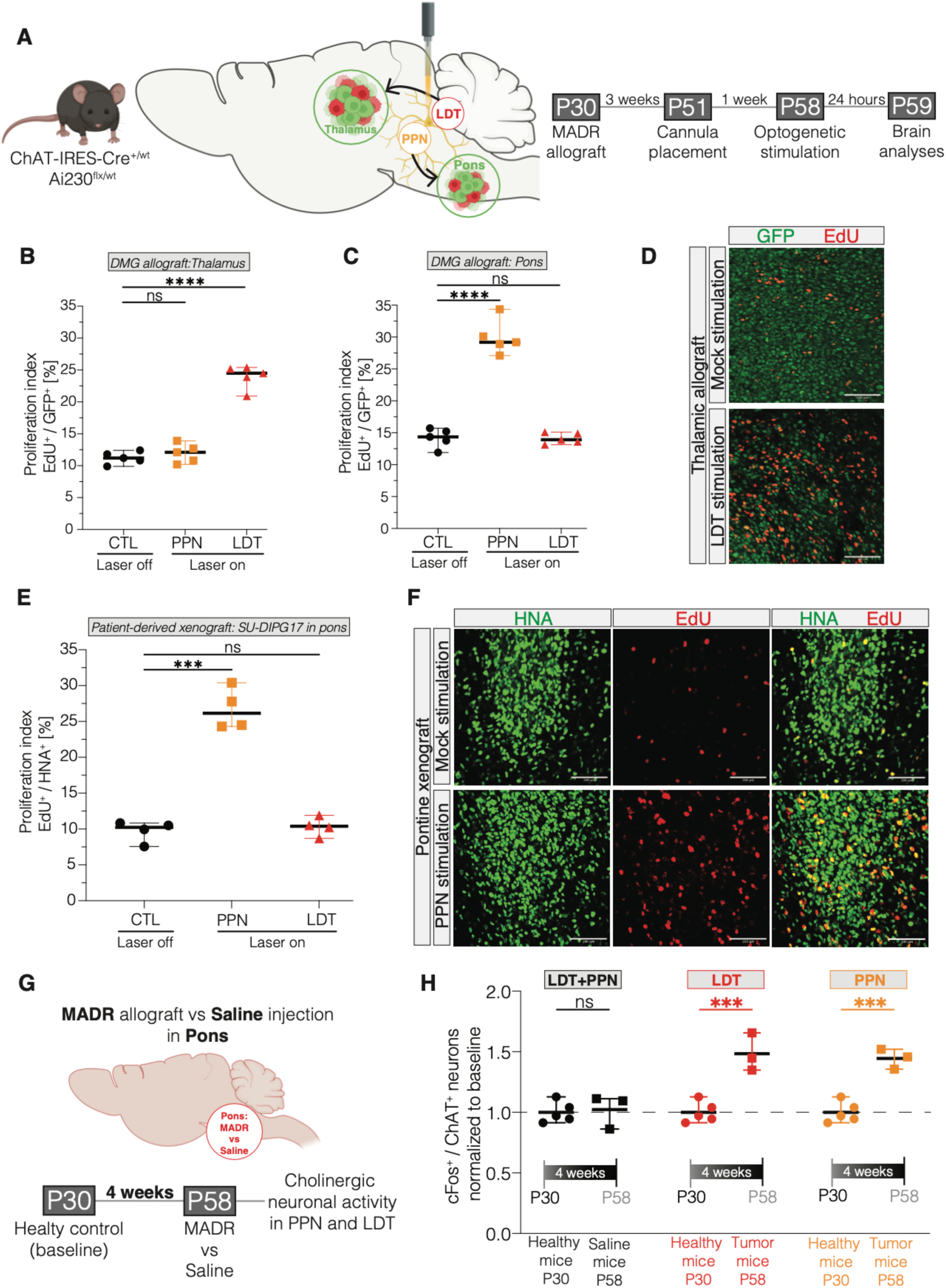
Interactions between cholinergic midbrain neurons and diffuse midline glioma. **(A)** Schematic of experimental paradigm for optogenetic stimulation of cholinergic neurons in either laterodorsal tegmentum nucleus (LDT) or pedunculopontine nucleus (PPN) in mice bearing H3K27M DMG. Four-week-old ChAT-IRES-Cre^+/wt^ x Ai230^flx/wt^ mice (P28-30) were allografted with a H3K27M MADR tumour model into either the pons or thalamus, with optic ferrule placement into the LDT or PPN three weeks after allografting. Optogenetic stimulation for 30 minutes of the LDT or PPN was performed four weeks after allografting, followed by perfusion 24 hours after stimulation. **(B)** Proliferation index (EdU^+^/GFP^+^) of thalamus allografts in mice either stimulated in PPN or LDT or non-stimulated (“CTL”) (CTL, PPN, and LDT, n=5 mice/group). Unpaired two-tailed t-test; ****p < 0.0001, ns: non-significant. Data shown as mean, error bars indicate range. **(C)** Proliferation index (EdU^+^/GFP^+^) of pons allografts in mice either stimulated in PPN or LDT or non-stimulated (“CTL”) (CTL, PPN, and LDT, n=5 mice/group). Unpaired two-tailed t-test; ****p < 0.0001, ns: non-significant. Data shown as mean, error bars indicate range. **(D)** Confocal micrographs showing proliferating GFP^+^ tumour cells in the right thalamus in non-stimulated (“CTL”) (upper images) and LDT-stimulated (“LDT”) (bottom images) mice. GFP: green, EdU: red, scale bars = 100μm. **(E)** Proliferation index (EdU^+^/HNA^+^) of patient-derived pontine xenografts (SU-DIPG 17) in mice either stimulated in PPN or LDT or non-stimulated (“CTL”) (CTL, PPN, and LDT, n=4 mice/group). Unpaired two-tailed t-test; ***p < 0.001, ns: non-significant. Data shown as mean, error bars indicate range. **(F)** Confocal micrographs illustrate proliferating HNA^+^ glioma cells in non-stimulated (“CTL”) (upper images) and PPN-stimulated (“PPN”) (bottom images) mice after xenografting patient-derived diffuse midline glioma line (SU-DIPG17) into pons. HNA: green, EdU: red, scale bars = 100μm. **(G)** Schematic of experimental paradigm for investigating reciprocal signalling effects of tumour cells to cholinergic midbrain neurons. Four-week-old ChAT-IRES-Cre^+/wt^ x Ai230^flx/wt^ mice (P28-30) were either allografted with a H3K27 DMG or injected with buffered saline (HBSS) into pons and were perfused four weeks after surgery. Cholinergic neuronal activity of LDT and PPN were measured by cFos staining and normalized to four-week-old healthy ChAT-IRES-Cre^+/wt^ x Ai230^flx/wt^ mice. **(H)** An increased neuronal activity of cholinergic neurons (cFos^+^ in ChAT^+^ neurons) in LDT and PPN was observed in tumour-bearing mice when compared to healthy and saline-injected mice (healthy, n=5 mice; saline, PPN, and LDT, n=3 mice). *Unpaired two-tailed t-test; *p < 0.05, **p < 0.01, ***p < 0.001, ns: non-significant*.

Patient-derived orthotopic xenograft models of DMG are complimentary to genetically engineered mouse models. We next tested the effects of cholinergic neuronal activity on patient-derived pontine H3.3-K27M DMG orthotopically xenografted to the pons. Viral vectors (AAV-hSyn-hChR2(H134R)::eYFP or AAV-hSyn::eYFP) were injected into the LDT or PPN of 4-week-old NOD-SCID-IL2-gamma chain-deficient (NSG) mice, followed by xenografting of patient-derived DMG cells (SU-DIPG-17) into the pons. Optogenetic stimulation was performed after 6 weeks of tumour engraftment, with perfusion conducted 24 hours later (**Supplementary Fig. 2A**). Consistent with the results above, stimulation of the PPN increased proliferation of patient-derived diffuse intrinsic pontine glioma/pontine diffuse midline glioma cells xenografted to the pons, whereas this effect was not observed after LDT stimulation (**Fig. 2E-F**). These findings mirror the observed OPC response *in vivo* and highlight cholinergic neuronal activity as a driver of DMG proliferation in a brain circuit-specific and midbrain nucleus-dependent manner.

Given previous studies demonstrating various activity-regulated proteins that promote glioma cell proliferation, we investigated the role of secreted factors in cholinergic activity-regulated DMG proliferation. Conditioned medium (CM) was collected from midbrain explants containing either PPN or LDT nuclei (**Supplementary Fig. 2B**). *Ex vivo* optogenetic stimulation of cholinergic neuronal cell bodies in either PPN or LDT explants generated CM that increased patient-derived H3K27M DMG (SU-DIPG-17) proliferation *in vitro* (**Supplementary Fig. 2C-D**). The conditioned medium was fractionated by molecular size, which revealed that the proliferative effect of the midbrain explant CM is attributable to macromolecules with a molecular weight between 10 to 100 kDa (**Supplementary Fig. 2E**), consistent with previous studies of activity-regulated paracrine factors that identified proteins neuroligin-3 (NLGN3)^4,5,9^ and BDNF^4,8,9^ as paracrine factors promoting glutamatergic neuronal activity-regulated glioma proliferation in cortex^4^ and optic nerve^9^. We found that ANA-12 (specific inhibitor of the BDNF-receptor TrkB) abolished the proliferative effect of cholinergic neuronal activity-regulated factors in PPN- or LDT-CM while the addition of Neurexin (to sequester NLGN3) only minimally decreased proliferation (**Supplementary Fig. 2F**). Concordant with the conclusion that BDNF is the chief paracrine growth factor released as a result of midbrain cholinergic neuronal activity in this experimental paradigm, elevated BDNF levels were found in LDT-CM (**Supplementary Fig. 2G**) and PPN-CM (**Supplementary Fig. 2H**), while NLGN3 was only mildly increased in midbrain cholinergic nuclei CM by western blot analysis (**Supplementary Fig. 2I**). Acetylcholine levels were mostly unchanged in midbrain explant CM following cholinergic neuronal activity (**Supplementary Fig. 2G-H**). These midbrain cholinergic nuclei explants contain cholinergic cell bodies but not axon terminals, which likely explains the lack of acetylcholine release into CM in this experimental paradigm. Taken together, these findings indicate that activity-regulated neurotrophin signaling from midbrain cholinergic neurons to TrkB on DMG cells contributes to the growth-promoting effects of cholinergic neuronal activity, at least locally in the midbrain.

### DMG cells increase cholinergic midbrain neuronal activity

Just as cortical and hippocampal neuronal activity can drive glioma proliferation and growth, glioma cells secrete paracrine factors that increase neuronal hyperexcitability in these regions^6,37,38^ and cause remodeling of functional neural circuits^39^. Such functional remodeling is evident in glioblastoma patients who exhibit whole-brain abnormalities and functional changes in brain areas distant from the tumor location^40–42^. This led us to hypothesize that midbrain cholinergic neurons may be reciprocally influenced by DMGs. We injected either H3K27M DMG cells (MADR model) or vehicle control (buffered saline, HBSS) into the pons of 4-week-old immunocompetent mice (ChAT-IRES-Cre^+/wt^ x Ai230^flx/wt^) and assessed cholinergic neuronal activity within both midbrain cholinergic nuclei after 4 weeks of tumor growth (**Fig. 2G**). Neuronal activity was measured by expression of the immediate early gene cFos in tumor- or saline vehicle control-injected mice and compared to the cFos expression of 4-week-old healthy littermate controls. Indeed, we observed increased midbrain cholinergic neuronal activity within both nuclei in DMG tumor-bearing mice, which was not observed in saline vehicle control-injected mice (**Fig. 2H**). These findings indicate bidirectional interactions between pontine DMG and cholinergic midbrain neurons whereby cholinergic neurons increase DMG growth and DMG cells increase cholinergic neuronal activity.

### Acetylcholine directly affects DMG growth and migration

To assess the effects of human cholinergic neurons on DMG, we co-cultured patient-derived DMG cells with cholinergic neurons derived from human induced pluripotent stem cells (hiPSCs). hiPSC-derived cholinergic neurons from a healthy 12-year-old male (**Fig. 3A**) were matched with a pontine DMG cell culture from an 8-year-old male (SU-DIPG-17) (**Fig. 3B**). Co-culture resulted in a two-fold increase in the glioma proliferation rate (**Fig. 3C**). hiPSC-derived cholinergic neurons and glioma cells formed a dense and extensive neuron-to-glioma network (**Fig. 3B, 3D**). Using a patient-derived DMG cell culture (SU-DIPG-13FL) expressing postsynaptic PSD95-RFP, cholinergic neuron-to-DMG synaptic structures were evident by pre- and post-synaptic puncta co-localization between presynaptic neurons and post-synaptic glioma cells within these networks (**Fig. 3D**).

**Figure 3.**
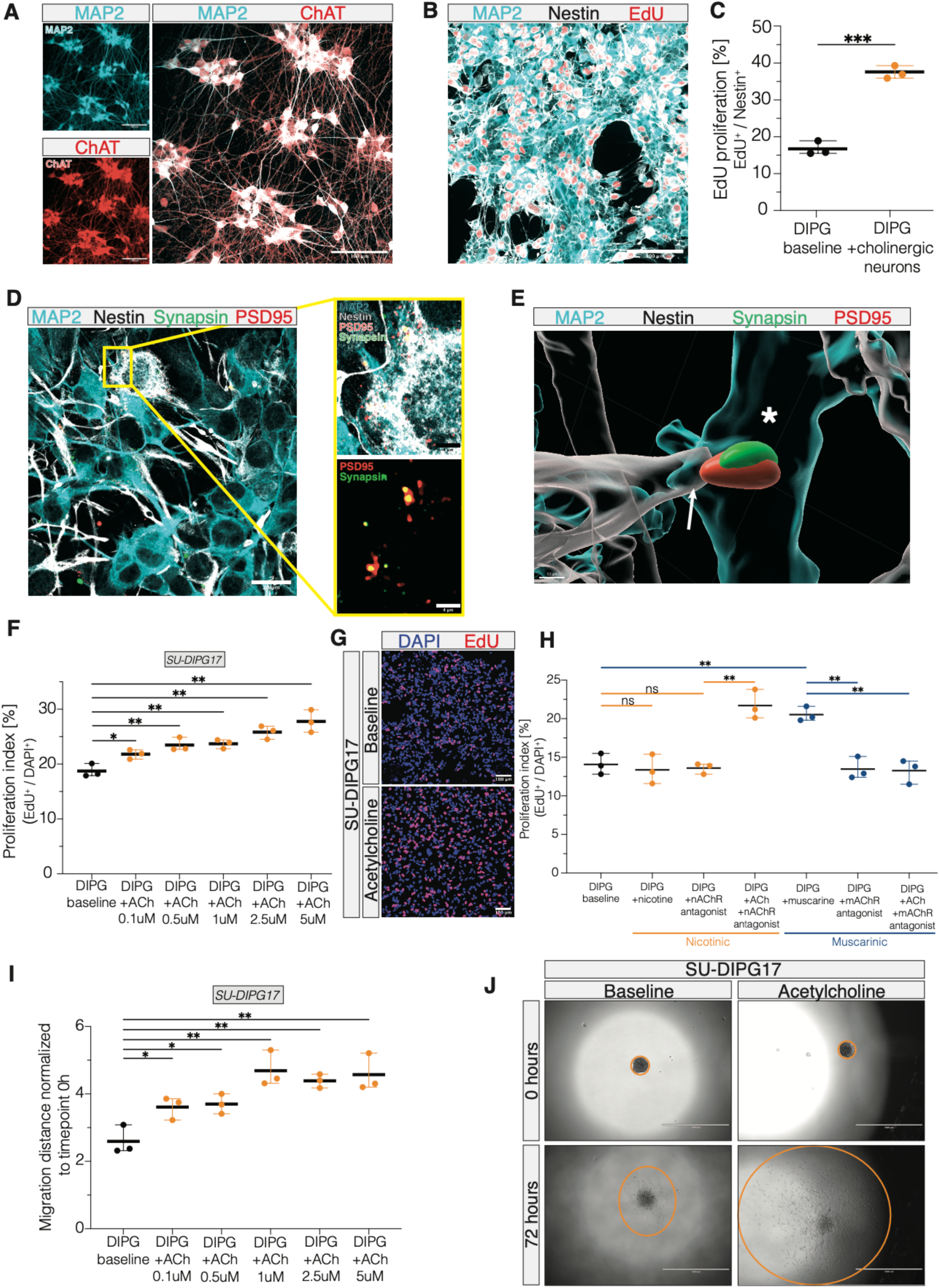
Direct effects of acetylcholine on diffuse midline glioma. **(A)** Confocal micrographs showing cholinergic neurons generated from human induced pluripotent stem cells (hiPSCs) of a healthy 12-year-old male. MAP2: turquoise, ChAT: red, merged: white, scale bars = 100μm. **(B)** Representative confocal micrographs demonstrating dense neuron-to-glioma networks after co-culturing hiPSC-derived cholinergic neurons and DMG cells. MAP2: turquoise, Nestin: white, EdU: red, scale bars = 100μm. **(C)** Quantification of glioma cell proliferation (EdU^+^/Nestin^+^/MAP2^-^) when co-cultured with hiPSC-derived cholinergic neurons. Unpaired two-tailed t-test; ***p < 0.001. Data shown as mean, error bars indicate range. **(D)** Confocal images of cholinergic neuron–glioma co-culture with PSD95–RFP-expressing DMG cells (SU-DIPG13FL-PSD95). Yellow boxes indicate colocalizations of synapsin (presynaptic cholinergic neuron) and PSD95 (postsynaptic glioma cell expressing PSD95-RFP). Nestin: white, MAP2: turquoise, PSD95-RFP: red, synapsin: green, scale bars = 20μm (left) and 4μm (right). **(E)** 3-dimensional rendering of the image in (D), illustrating presynaptic cholinergic neuron (MAP2, turquoise, asterisk) with presynaptic puncta (green, synapsin) co-localizing with postsynaptic puncta (PSD95-RFP, red) expressed by postsynaptic glioma cell (nestin, white, arrow). **(F)** Proliferation index (EdU^+^/DAPI^+^) of a patient-derived DMG cell line (SU-DIPG17) after exposure to different concentrations of acetylcholine. **(G)** Representative confocal micrographs showing the proliferation of a patient-derived cell line without acetylcholine (upper images) and after exposure to 5μM acetylcholine (bottom images). DAPI: blue, EdU: red, scale bars = 100μm. **(H)** Proliferation index (EdU^+^/DAPI^+^) of a patient-derived cell line (SU-DIPG17) after exposure to various muscarinic and nicotinic receptor agonists (nicotine and muscarine) and antagonists (mecamylamine and scopolamine). One-way analysis of variance (ANOVA) with Tukey’s post hoc analysis; **p < 0.01, ns: non-significant. Data shown as mean, error bars indicate range. **(I)** 3D migration assay analysis comparing distance of glioma cell spread 72 h after seeding after exposure to acetylcholine. One-way analysis of variance (ANOVA) with Tukey’s post hoc analysis; *p < 0.05, **p < 0.01. Data shown as mean, error bars indicate range. **(J)** Representative images showing the glioma cell migration at timepoint zero and after 72h in control- and acetylcholine-treated wells. Scale bars = 1000μm.

The tumor growth-promoting effects of cholinergic neurons *in vivo* and *in vitro* suggest that acetylcholine release at axon terminals may directly act on DMG cells. To assess the direct effects of acetylcholine on DMG cells, we tested the proliferation rate of patient-derived H3K27M DMG cell cultures (n = 4 distinct patient-derived DMG cultures, **Supplementary Table 1**) exposed to varying concentrations of acetylcholine *in vitro*. These experiments revealed a dose-dependent increase in DMG cell proliferation following exposure to acetylcholine, which plateaued at higher concentrations (**Fig. 3E-F** and **Supplementary Fig. 3A-C**). Muscarinic receptor blockers abrogated the effects of acetylcholine, while nicotinic receptor blockers had no effect on the proliferation caused by acetylcholine (**Fig. 3G**). Neither muscarinic nor nicotinic receptor blockers affected glioma cell proliferation in the absence of acetylcholine (**Fig. 3G**). Furthermore, muscarine (a direct muscarinic receptor agonist) induced glioma cell proliferation to the same extent as acetylcholine, while nicotine (a direct nicotinic receptor agonist) exerted no effect (**Fig. 3G**). Taken together, these data indicate that the proliferative effect of acetylcholine is mediated by muscarinic receptors rather than nicotinic receptors.

Glutamatergic and GABAergic neurons promote the proliferation and progression of H3K27M DMG in part through synaptic signaling^4,8,28^. Given the neuromodulatory functions of acetylcholine^43^, we next asked whether acetylcholine alters the known^6,8^ proliferative effect of cortical neuron co-culture on glioma cells. Early postnatal mouse pup (P0-P1) cortical preparation generated mixed glutamatergic (Vglut^+^) and GABAergic (GAD65^+^) cortical neuron cultures (**Supplementary Fig. 3D**). As expected^6,8^, cortical neuron-DMG co-culture markedly increased the proliferation rate of DMG cells, tested in two independent patient-derived cultures (SU-DIPG-17 and SU-DIPG-92; **Supplementary Fig. 3E-G**). Addition of acetylcholine substantially augmented this effect, further increasing the glioma cell proliferation rate in a dose-dependent manner (**Supplementary Fig. 3F-G**). Future work will be required to determine if this effect of acetylcholine in the context of cortical neuron co-culture simply represents an additive effect of acetylcholine-induced glioma proliferation to the proliferation-inducing effects of cortical neurons on DMG cells, or if acetylcholine is also acting as a neuromodulator to augment glutamatergic^6^ and/or GABA-ergic^28^ synaptic signaling in DMG cells.

Previous studies indicate that neuronal activity promotes the widespread dissemination of glioma cells^29,30^. Concordantly, we found that acetylcholine exposure increased the migration of DMG cells *in vitro* in a dose-dependent manner (**Fig. 3H-I**). Taken together, these findings highlight a direct effect of acetylcholine on proliferation and infiltration in DMG, representing a second mechanism - in addition to the cholinergic neuronal activity-regulated neurotrophin signaling described above – by which cholinergic neuronal activity modulates DMG pathophysiology.

### CHRM1 and CHRM3 are therapeutic targets in DMG

The proliferation-promoting effects of acetylcholine *in vitro* and cholinergic long-range projections *in vivo* raise questions about the cholinergic receptor profile of DMG cells. We further tested receptor gene expression in multiple single-cell and single-nucleus RNA sequencing (sc/snRNAseq) datasets from primary patient tumor samples across various central nervous system tumors, including 54 DMG samples. Cholinergic receptor genes exhibited varying levels of expression across malignant cells within DMG tumors (**Supplementary Fig. 4A)**, reflecting intratumoral heterogeneity - a phenomenon well-described in this disease^12^ and in gliomas in general^44–46^. To highlight intertumoral heterogeneity we compared pseudo-bulk expression in malignant cells across samples (**Fig. 4A**). Notably, the muscarinic receptor *CHRM3* was found to be highly expressed across all glioma subtypes, ependymoma, and medulloblastoma, but not in neuroendocrine pituitary tumors (**Fig. 4A**). We next sought to determine how intratumor heterogeneity and cell states are correlated to cholinergic receptor gene expression. We initially focused on the OPC-like cell state, which reflects the cancer stem cell/tumor-initiating cell subpopulation^13^ and has been observed to be particularly prevalent in DMG tumors compared to other glioma subtypes^44^. Examining differentially expressed genes associated with either muscarinic or nicotinic receptor genes across all tumor samples revealed a specific association between muscarinic receptor genes and *PDGFRA* – a hallmark gene for OPCs - exclusively in DMG (**Supplementary Table 2**). We further tested correlations between cholinergic receptor gene expression and OPC scores, both within tumor cells and across samples at the pseudo-bulk level. We found that the M1 muscarinic receptor gene *CHRM1* exhibited the highest values on both measures (**Fig. 4B**), stronger than but similar to the values for the M3 muscarinic receptor gene *CHRM3* (**Fig. 4B**). Both *CHRM1* and *CHRM3* expression are enriched in the OPC-like cell state of DMG (**Fig. 4C-D)**. Notably, in medulloblastoma samples, which is an embryonal, non-glioma tumor entity, we observed no correlation between any cholinergic receptor gene and an OPC score (**Supplementary Fig. 4B**). Further comparisons of intertumoral correlations of cholinergic receptor gene expression across various tumor types and with different cell states underscored the association between *CHRM1* and OPC-like states in DMG using the Neftel et al^44^ and a pan-cancer study-derived cell states^47^ as references (**Supplementary Fig. 4C**). While not the most highly expressed cholinergic receptor gene in glioma, *CHRM1* seems to play a unique role in OPC-like DMG cells, distinguishing it from other CNS tumor entities such as glioblastoma, ependymoma, and medulloblastoma. By contrast, CHRM3 appears to be important in multiple other tumors, including glioblastoma^48,49^.

**Figure 4.**
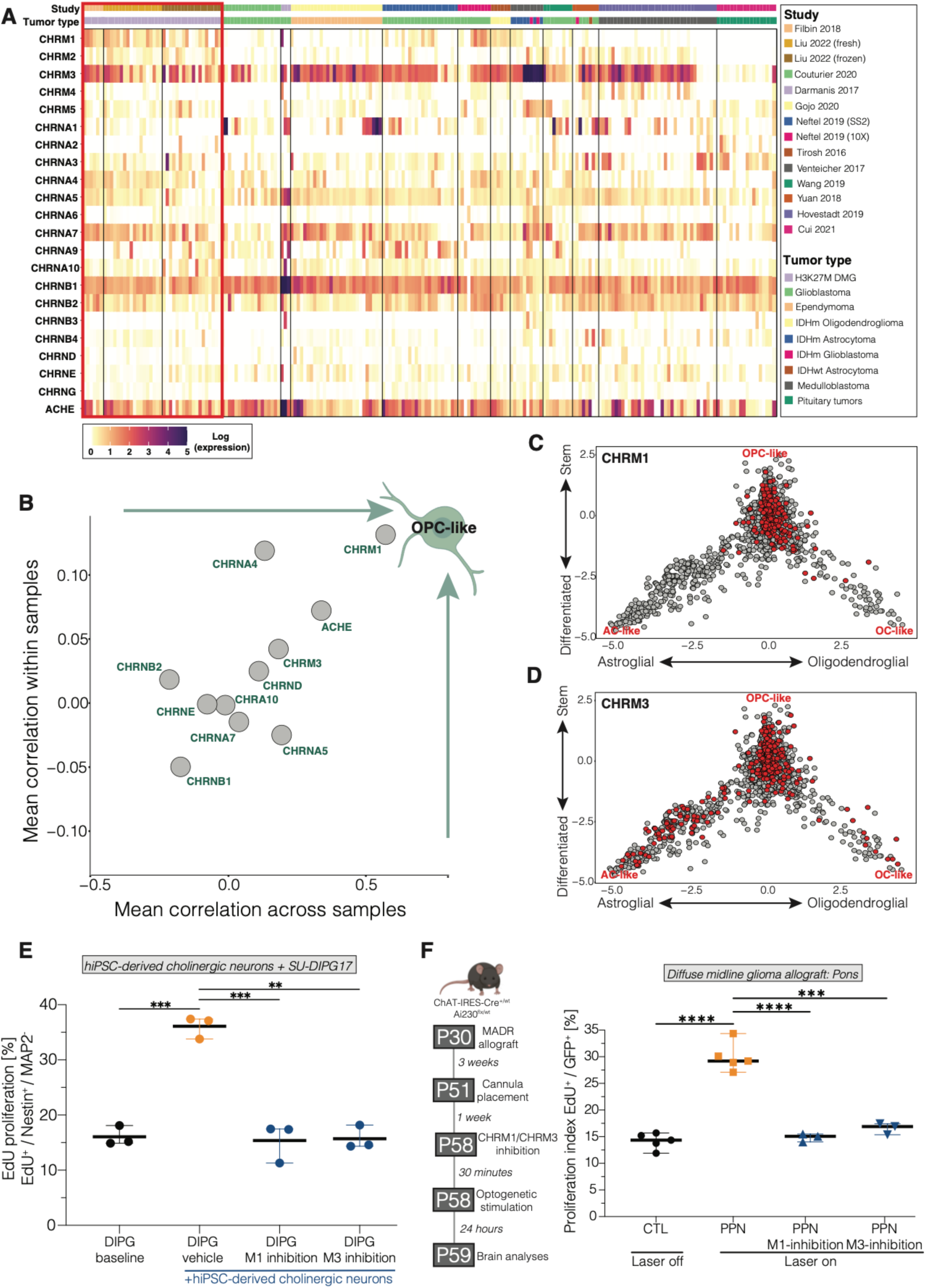
Cholinergic receptor gene expression and receptor mechanisms in gliomas. **(A)** Heatmap of pseudo-bulk analysis of cholinergic receptors gene expression across central nervous system tumours from various studies with single-cell or single-nucleus RNA sequencing data. The red-marked boxes indicate studies with DMG samples. **(B)** Scatter plot correlating cholinergic receptor gene expression values with an OPC-like score in DMG samples. **(C)** Two-dimensional representation of the association between CHRM1 expression (red dots: centered value > 1) and the OPC-like (y axis) as well as OC-like and AC-like (x axis) scores for H3K27M DMGs. **(D)** Two-dimensional representation of the association between CHRM3 expression (red dots: centered value > 1) and the OPC-like (y axis) as well as OC-like and AC-like (x axis) scores for H3K27M DMGs. **(E)** Proliferation index (EdU^+^/Nestin^+^) of a patient-derived DMG cell line (SU-DIPG17) when co-cultured with hiPSC-derived cholinergic neurons and treated with M1 (VU0255035) and M3 (4-DAMP) receptor antagonists. Unpaired two-tailed t-test; **p < 0.01, ***p < 0.001. Data shown as mean, error bars indicate range. **(F)** Experimental paradigm (left) and proliferation index (EdU^+^/GFP^+^) of pontine allografts in mice optogenetically stimulated in PPN or non-stimulated controls (“CTL”), with or without administration of M1 (VU0255035) or M3 receptors (4-DAMP) pharmacological inhibitors prior to optogenetic stimulation of cholinergic neurons in mice bearing H3K27M DMG (MADR model) in the pons. The mice who did not receive M1 or M3 blockers were administered vehicle control. (n=4 mice/group in control and PPN-stimulated groups; n=3 mice/group in PPN-stimulated mice treated with M1-antagonist or M3-antagonist). The control group received vehicle control. Unpaired two-tailed t-test; ***p < 0.001, ****p < 0.0001. Data shown as mean, error bars indicate range.

Since *CHRM1* and *CHRM3* both appear to play significant roles in DMG, as revealed by their potential as therapeutic targets, we tested their role in DMG proliferation *in vitro* and *in vivo.* Utilizing the iPSC-derived cholinergic neuron co-culture model, we found that antagonists to M1 receptors (VU0255035) and M3 receptors (4-DAMP) blocked the proliferation-inducing effects of cholinergic neurons on DMG (**Fig. 4E**) and the number of cholinergic neuron-to-glioma puncta colocalizations (**Supplementary Fig. 4D**). We next tested the effects of cholinergic antagonists on DMG tumor proliferation in vivo. Mice bearing pontine DMG allografts were treated with M1 antagonists, M3 antagonists, or vehicle control prior to optogenetic stimulation or identical manipulation (surgery, handling, etc.) without light. We found that the administration of either M1 or M3 antagonist completely abolished the proliferative effect of PPN cholinergic neuronal activity on pontine DMG *in vivo* (**Fig. 4F, Supplementary Fig. 4E**). These findings highlight the importance of muscarinic receptors M1 and M3 in activity-regulated cholinergic neuron-to-glioma interactions.

## DISCUSSION

H3K27M-altered diffuse midline gliomas are aggressive brain cancers that occur in midline structures, chiefly the pons (also called DIPG), thalamus and spinal cord. Both DMGs and OPCs – the DMG cell of origin^12–17^ - proliferate in response to the activity of glutamatergic neurons^4,23^. Here, we have tested the effects on normal OPCs and malignant OPC-like tumor cells of cholinergic neurons located in the midbrain, a structure located between the pons and thalamus that is frequently invaded by both pontine and thalamic DMGs. The axon projections of cholinergic neurons in the two distinct cholinergic nuclei of the midbrain - LDT and PPN - project robustly to the thalamus and pons, respectively. We found that both normal OPCs and DMG cells in pons and thalamus proliferate in response to cholinergic neuronal activity in a circuit-specific manner, with acetylcholine acting on muscarinic M1 and M3 acetylcholine receptors enriched in the OPC-like DMG cellular subpopulation.

Given the oligodendroglial lineage precursor cellular origins of DMGs and the similarities between normal OPC and DMG cells^12–17^, it is useful to study both in parallel (for review, see^18^). Like glioma cells^4^, glutamatergic neuronal activity promotes the proliferation of normal OPCs^23^, as well as oligodendrogenesis and adaptive, activity-regulated myelin changes that tune neural circuit function^23,26,50^; but the possible role of cholinergic neurons in myelin plasticity is not yet clear. Here we found the that cholinergic long-range projections similarly influence both normal and malignant cell proliferation. However, the effects of cholinergic neuronal activity on the generation of new oligodendrocytes and on myelination – of cholinergic axons or other axons – remains to be elucidated. OPC proliferation can reflect either the generation of new oligodendrocytes, or a blockade in differentiation that maintains the precursor in a proliferating state. Cholinergic signaling in OPCs is complex: in cultured OPCs, muscarinic signaling promotes OPC proliferation^51,52^, increased survival^53^, and prevents differentiation^54^, while nicotinic signaling promotes OPC differentiation into mature oligodendrocytes^55^. Thus, acetylcholine acts through muscarinic signaling to block oligodendrogenesis while acting through nicotinic signaling to promote oligodendrogenesis. Concordantly, muscarinic blockade promotes remyelination^54,56^ after demyelinating injury, and muscarinic antagonists like clemastine – an anti-histamine drug with anticholinergic properties - have been^57^ and continue to be tested in clinical trials for multiple sclerosis (NCT02521311, NCT05338450). We find that cholinergic neuronal activity promotes OPC proliferation in midline brain regions and in cortex, but not in other regions like the hippocampus and VTA. OPCs are regionally and temporally heterogeneous^58^, and whether different OPC populations express varying proportions of nicotinic to muscarinic receptors to account for these differences is an open question that may help to explain the regional heterogeneity in functional responses observed here. It will furthermore be elucidating if regional heterogeneity in therapeutic responses is uncovered during these promising remyelination clinical studies^57^ (NCT02521311, NCT05338450).

The tumor growth-promoting effects of cholinergic neuronal activity on DMG cells are mediated by both acetylcholine and BDNF release, although these two mechanisms are spatially enriched at different locations. The proliferative effects of long-range cholinergic neuron projections to thalamus and pons are chiefly mediated through acetylcholine signaling via M1 and M3 receptors, as the pro-tumor effects of cholinergic neuronal activity are blocked by M1 or M3 antagonists *in vitro* and *in vivo*. Furthermore, cholinergic neurons influence DMG cells through BDNF-TrkB signaling, which is known to promote proliferation and malignant synaptic plasticity in gliomas^4,8^. BDNF release appears to occur closer to the neuronal soma, as explants of cholinergic midbrain nuclei containing neuronal soma and local neuronal projections such as short-range axons and dendrites - but not longer axons - exhibited activity-regulated BDNF release that promoted DMG proliferation. The extent to which DMG cells in the midbrain, a site commonly invaded by DMGs originating either in the pons or thalamus, are influenced by cholinergic neuron-derived BDNF remains to be tested; BDNF can be released by numerous neuron types, and inhibition of the BDNF receptor TrkB reduces DMG growth and extends survival in xenograft mouse models^8^.

Neuronal activity robustly drives the progression of DMG and other glioma^8–10,30,39^. These effects are mediated through both paracrine^4,10,30^ and synaptic^6–8,29^ neuron-to-glioma interactions. Here, we uncovered a role for acetylcholine, which can function as a paracrine growth factor in DMG, but also may be signalling at cholinergic neuron-to-glioma synapses. Concordant with recent evidence that cholinergic neurons can form synapses with glioblastoma cells^48,49^, we found evidence of cholinergic neuron-to-DMG cell synaptic structures. The extent to which cholinergic synaptic signaling, or cholinergic modulation of glutamatergic synaptic signaling, accounts for the glioma growth-promoting effects of acetylcholine will be important areas for future study.

Interactions between neurons and glioma are bidirectional, and glioma cells can profoundly affect neuronal function by increasing neuronal excitability^37,38,59–62^ and functionally remodeling neural circuits^39^. We observed an effect of DMG on the baseline neuronal activity of cholinergic midbrain neurons. Given the important neuromodulatory effects of cholinergic signaling on cognition and other brain functions (for review, see^43^), this raises the possibility that tumor-induced dysregulation of cholinergic neurons – and possibly other neuromodulatory neuron types - in the brainstem may contribute to the cognitive, behavioral and emotional symptoms that DMG patients frequently experience^63^. Further work on interactions between neuromodulatory brainstem neurons and DMG tumor cells is needed.

Integration of primary tumor single-cell/single-nucleus datasets revealed the muscarinic receptor CHRM3 as the most highly expressed cholinergic receptor gene across various CNS tumours, including DMG. Further detailed analysis identified CHRM1 as a unique target, upregulated solely in the OPC-like cell state within DMG. Our *in vitro* and *in vivo* findings highlight the potential of these two receptors as therapeutic targets for DMG. The nervous system regulates a wide range of cancers throughout the body (for reviews, please see^2,3,64^), and emerging principles in the burgeoning field of Cancer Neuroscience are becoming evident. Cholinergic signaling through muscarinic receptors is required for glandular organogenesis^65^ and has emerged as a frequent mechanism regulating tumor pathophysiology, evident not only in brain tumors as explored here, but also in diverse cancer types including gastric^66^, colon^67^ and prostate cancers^68^. Blocking M1 or M3 muscarinic receptors using tool compounds demonstrate the therapeutic promise of targeted muscarinic receptor inhibition for DMG. Beyond DMG, the literature indicates broad potential applications for clinically development of specific M1 and/or M3 inhibitors to target muscarinic signaling in cancer.

Taken together, the results presented here implicate cholinergic brainstem neurons as an important driver of diffuse midline glioma pathophysiology. Each neuron type studied to date Fihas proven to contribute to DMG growth and progression through targetable mechanisms. A comprehensive understanding of the neuroscience of diffuse midline gliomas will enable the development of effective combination therapy strategies for these lethal brain cancers of childhood.

**Supplementary Figure 1.**
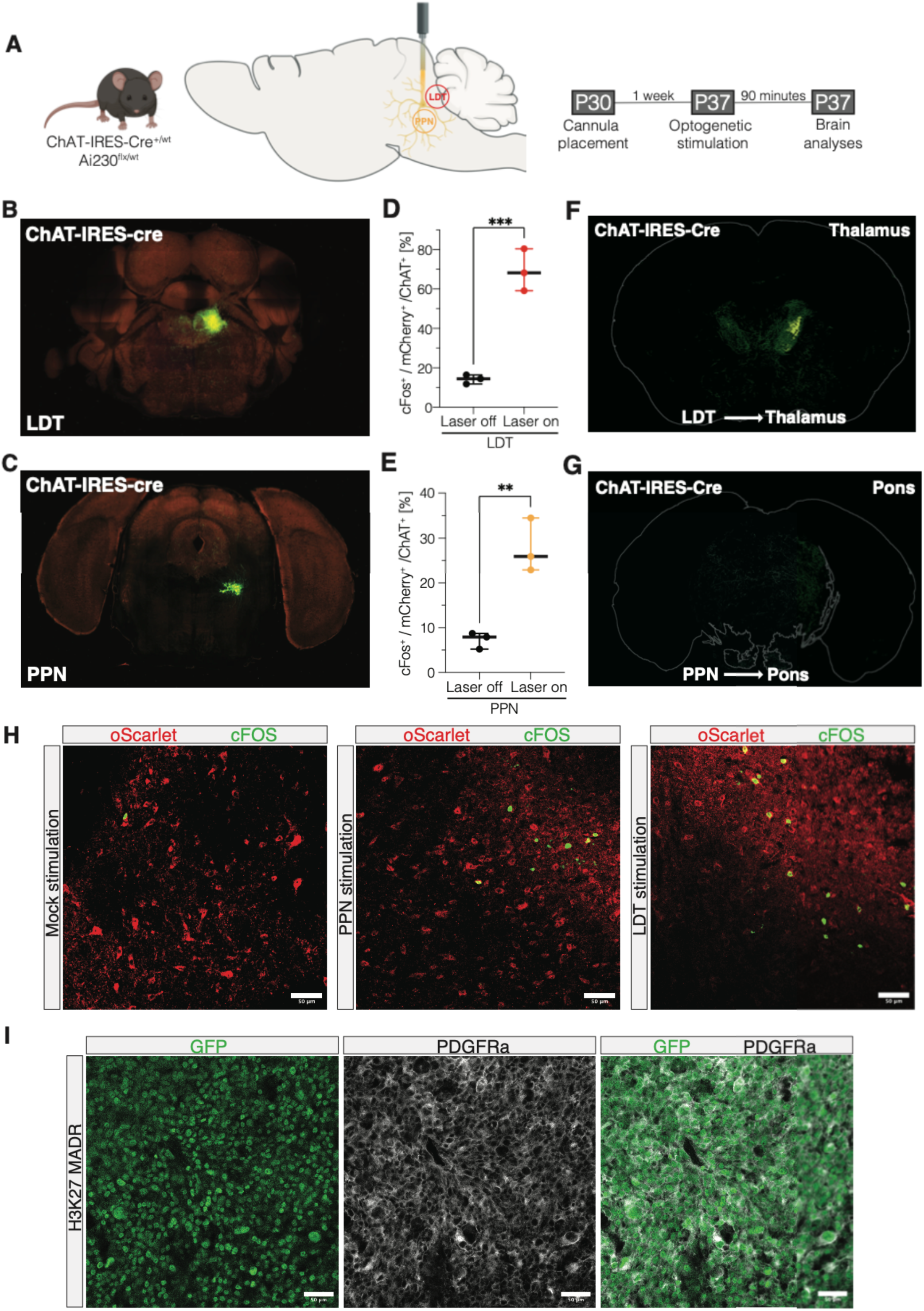
Validation of experimental paradigms. **(A)** Schematic of experimental paradigm for validation of optogenetic stimulation of cholinergic neurons in either laterodorsal tegmentum nucleus (LDT) or pedunculopontine nucleus (PPN). 5-week-old ChAT-IRES-Cre^+/wt^ x Ai230^flx/wt^ mice (P35-38) were stimulated for 30 minutes followed by perfusion after 90 minutes to assess neuronal activity changes by cFos staining. **(B)** – **(C)** Cre-dependent AAV tracing of axonal projections within the injection site in the **(B)** LDT and **(C)** PPN. Images are obtained from the Allen Mouse Brain Connectivity Atlas (connectivity.brain-map.org/projection/experiment/264566672)^69–72^. **(D)** Quantification of activated cholinergic neurons (cFos^+^/mCherry^+^/ChAT^+^) after optogenetic stimulation of LDT (CTL, LDT, n=3 mice). Unpaired two-tailed t-test; ***p < 0.001. Data shown as mean, error bars indicate range. **(E)** Quantification of activated cholinergic neurons (cFos^+^/mCherry^+^/ChAT^+^) after optogenetic stimulation of PPN (CTL, PPN, n=3 mice). Unpaired two-tailed t-test; **p < 0.01. Data shown as mean, error bars indicate range. **(F) - (G)** Cre-dependent AAV tracing of axonal projections from an injection site in the LDT and PPN. Ipsilateral projection targets include the thalamus for LDT (**F**) and pons for PPN (**G**). Images are obtained from the Allen Mouse Brain Connectivity Atlas (connectivity.brain-map.org/projection/experiment/264566672)^69–72^. **(H)** Confocal micrographs showing cFos staining in cholinergic neurons of non-stimulated mice (left image), PPN-stimulated mice (middle image), and LDT-stimulated mice (right image). ChRmine-oScarlet: red, cFos: green, scale bars = 50μm. **(I)** Confocal micrographs highlighting the OPC-like character (Pdgfra^+^) in tumour cells (GFP^+^) of the H3K27M MADR model. GFP: green, Pdgfra: white, scale bars = 50μm. *Related to Figures 1 and 2.*

**Supplementary Figure 2.**
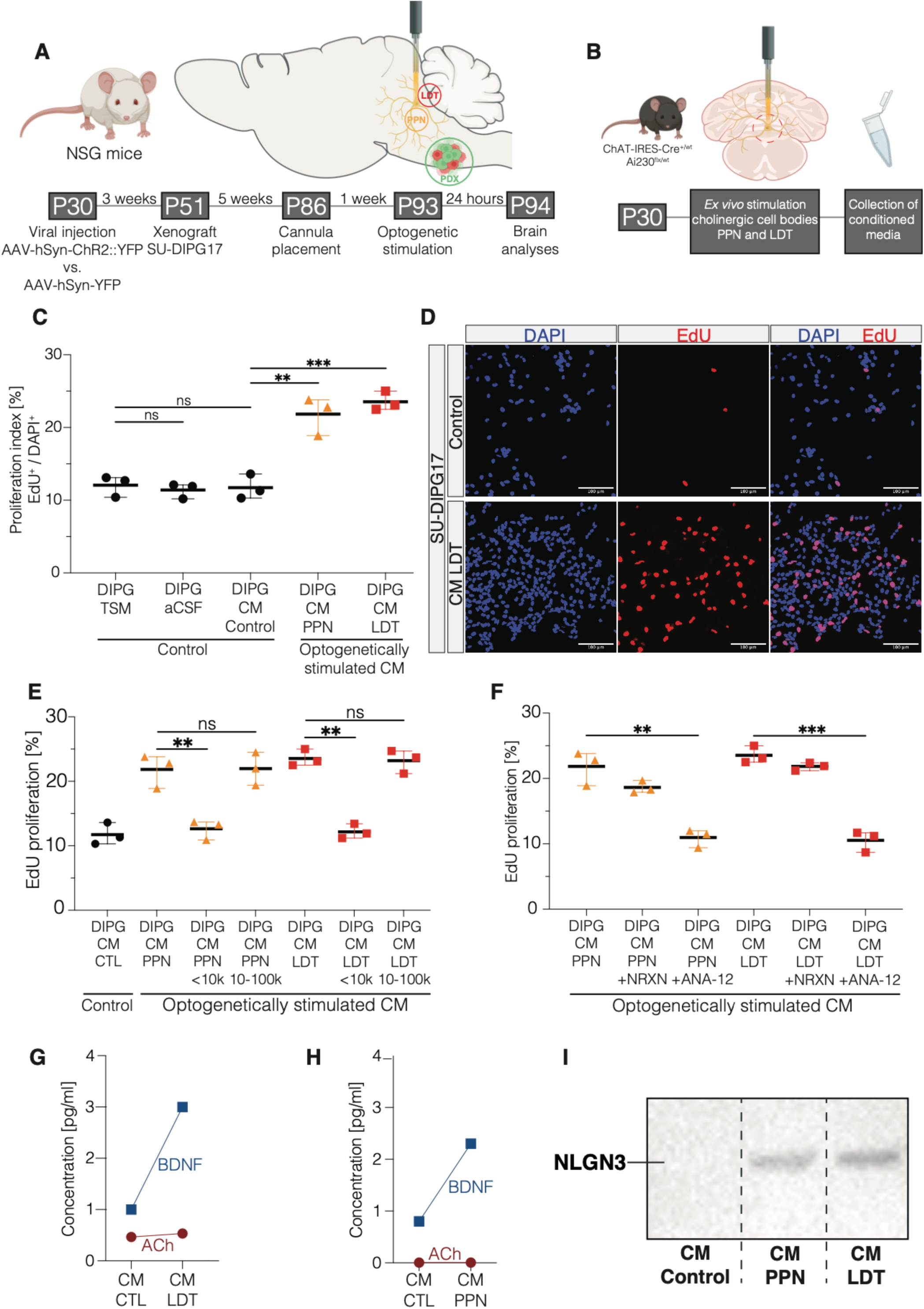
Midbrain cholinergic neurons promote DMG growth in part through paracrine factors such as BDNF. **(A)** Schematic of experimental paradigm for xenografting with optogenetic stimulation of either laterodorsal tegmentum nucleus (LDT) or pedunculopontine nucleus (PPN) in immunodeficient mice. Four-week-old NSG mice (P28-30) were injected with AAV-hSyn-ChR2(H134R)::eYFP or AAV-hSyn::eYFP into LDT or PPN and xenografted into the pons with patient-derived H3K27-altered DIPG cells (SU-DIPG17) three weeks after viral vector delivery. After five weeks of glioma growth, optical ferrules were placed either in the LDT or PPN and optogenetic stimulation was performed one week later. Mice were perfused 24 hours after optogenetic stimulation. **(B)** Schematic of experimental paradigm for collection of conditioned media (CM) after *ex vivo* optogenetic stimulation of cholinergic neuronal cell bodies within the LDT or PPN in midbrain explants of 4-week-old ChAT-IRES-Cre^+/wt^ x Ai230^flx/wt^ mice. **(C)** Quantification of DMG cell proliferation (EdU^+^/DAPI^+^) when adding CM after *ex vivo* stimulation of LDT or PPN compared to CM from non-stimulated slices. One-way analysis of variance (ANOVA) with Tukey’s post hoc analysis; **p < 0.01, ***p < 0.001, ns: non-significant. Data shown as mean, error bars indicate range. **(D)** Confocal micrographs showing EdU-labeled DMG proliferation after adding CM from non-stimulated slices (upper images) and from LDT stimulation (bottom images). DAPI: blue, EdU: red, scale bars = 100 μm. **(E)** Proliferation index (EdU^+^/DAPI^+^) after fractionation of the CM by molecular weight using size-exclusion ultracentrifugal filters on CM from LDT and PPN expalnts. One-way analysis of variance (ANOVA) with Tukey’s post hoc analysis; **p < 0.01, ns: non-significant. Data shown as mean, error bars indicate range. **(F)** Quantification of glioma cell proliferation (EdU^+^/DAPI^+^) with addition of neurexin (which binds to and sequesters NLGN3^4^) or ANA-12 (specific TrkB-inhibitor) to the CM from LDT and PPN. One-way analysis of variance (ANOVA) with Tukey’s post hoc analysis; **p < 0.01, ***p < 0.001. Data shown as mean, error bars indicate range. **(G) – (H)** Measurement of BDNF and acetylcholine levels in CM from (G) LDT, and (H) PPN. **(I)** Western blot analysis of NLGN3 in CM from non-stimulated midbrain explants (“Control”) as well as CM from optogenetically stimulated PPN explants and from LDT explants. *Related to Figure 2.*

**Supplementary Figure 3.**
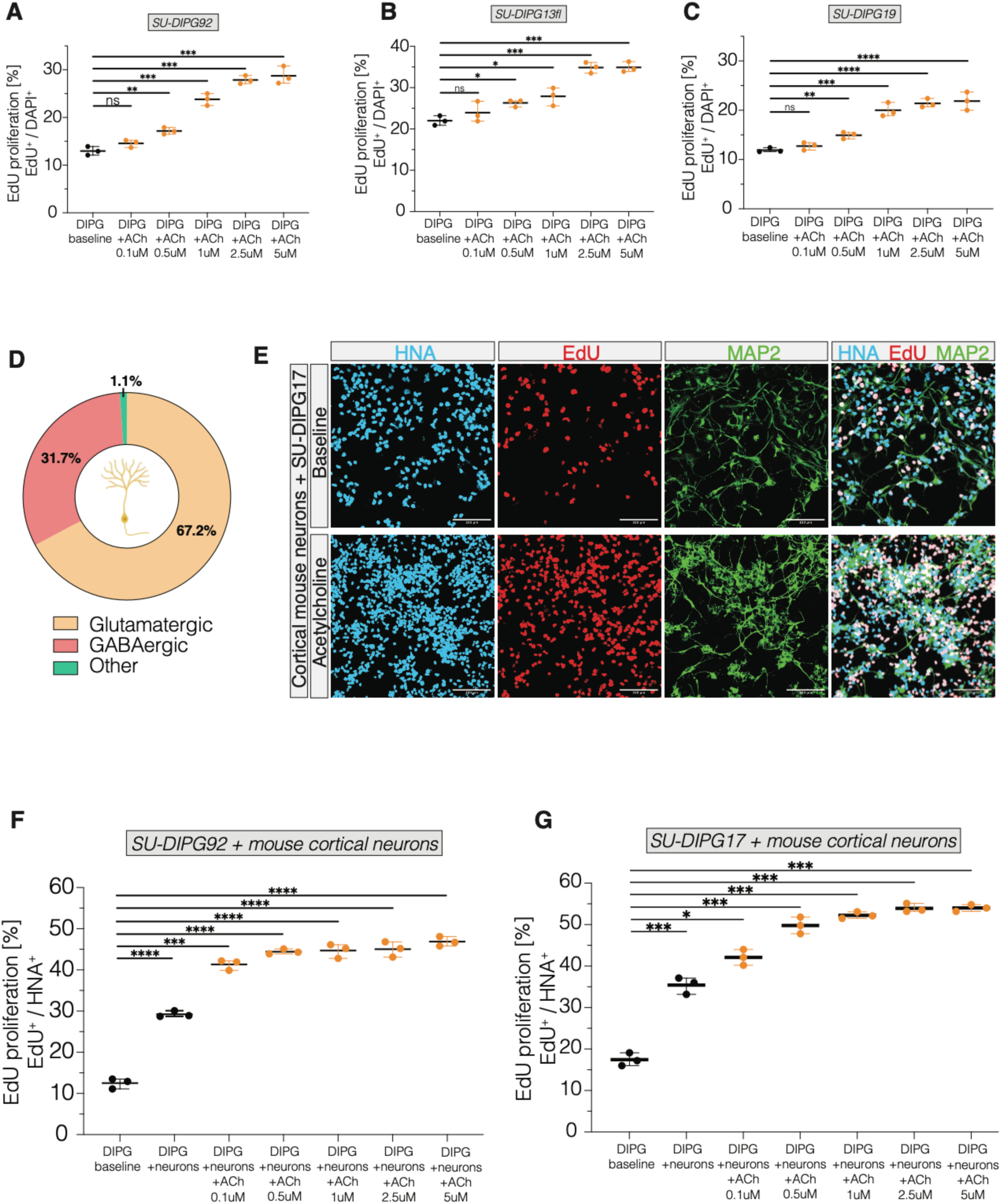
Acetylcholine promotes DMG cell proliferation in monoculture and cortical neuron co-culture in a dose dependent manner. **(A) – (C)** Proliferation index (EdU^+^/DAPI^+^) of three patient-derived H3K27M DMG cell cultures (**A:** SU-DIPG92, **B:** SU-DIPG13fl, **C:** SU-DIPG19) after exposure to different concentrations of acetylcholine. One-way analysis of variance (ANOVA) with Tukey’s post hoc analysis; *p < 0.05, **p < 0.01, ***p < 0.001, ****p < 0.0001, ns: non-significant. Data shown as mean, error bars indicate range. **(D)** Proportions of neuronal subpopulations within the cortical neuron cultures from mouse pups used for the co-culture experiment shown in Supplementary Fig. 3E-G. **(E)** Confocal micrographs representing the proliferative effect of adding acetylcholine (1μM) to a neuron-glioma co-culture. HNA: blue, EdU: red, MAP2: green, scale bars = 100μm. **(F) – (G)** Proliferation index (EdU^+^/HNA^+^) of patient-derived DMG cell cultures when co-cultured with cortical neurons from mouse pups and after exposure to different concentrations of acetylcholine. One-way analysis of variance (ANOVA) with Tukey’s post hoc analysis; p < 0.05, ***p < 0.001, ****p < 0.0001. Data shown as mean, error bars indicate range. *Related to Figure 3.*

**Supplementary Figure 4.**
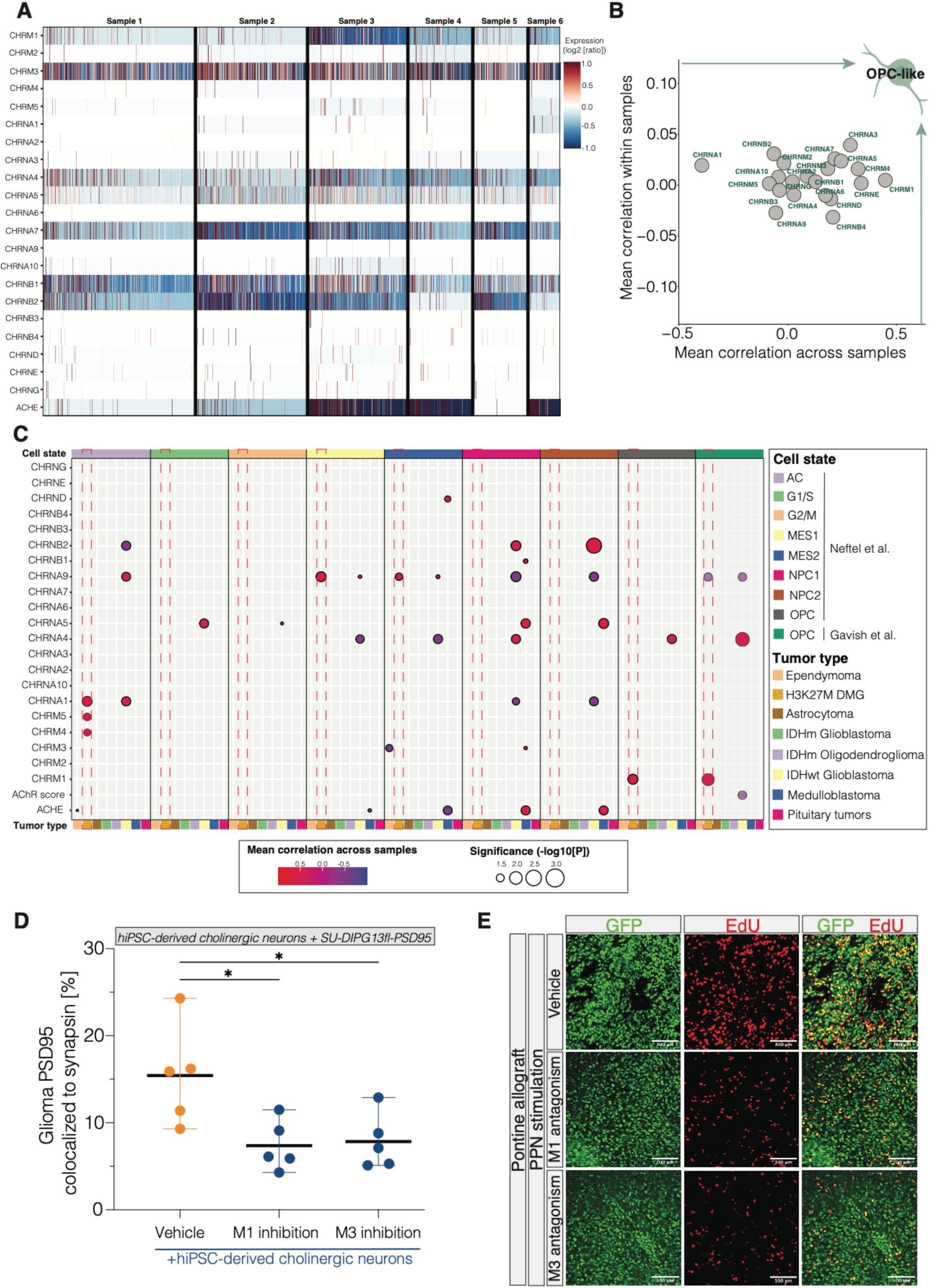
Muscarinic targets for DMG therapy. **(A)** Heatmap illustrating intra- and intertumoral heterogeneity of acetylcholine receptor gene expression in six scRNA-seq DMG samples (Filbin dataset^13^). The data were centered across all cells per sample. **(B)** Scatter plot correlating cholinergic receptor gene expression values with an OPC-like score in medulloblastoma samples. **(C)** Correlation of cholinergic receptor gene expressions with different cell like-states (AC, G1/S, G2/M, MES-1, MES-2, NPC-1, NPC-2, and OPC) in all available CNS tumour entities. The radius of each circle corresponds to the −log^10^(p-value), and the circle filling is the correlation. Non-significant p-values are not shown. Red dashed boxes indicate studies containing DMG samples. **(D)** Colocalization of PSD95 (postsynaptic glioma cell) and synapsin (presynaptic cholinergic neuron) in a PSD95-labelled DMG cell line when co-cultured with hiPSC-derived cholinergic neurons and treated with M1 (VU0255035) and M3 (4-DAMP) receptor antagonists. Unpaired two-tailed t-test; *p < 0.05. Data shown as mean, error bars indicate range. **(E)** Confocal micrographs showing proliferating GFP^+^ tumour cells in the pons in PPN-stimulated mice without drug administration (upper row), M1 receptor antagonism (middle row) or M3 (bottom row) receptor antagonism. GFP: green, EdU: red, scale bars = 100μm. *Related to Figure 4.*

## METHODS

### Patient-Derived Diffuse Midline Glioma Cells

Diffuse midline glioma cultures were established from patient-derived specimens with informed consent, following a protocol approved by the Stanford University Institutional Review Board (IRB). The utilized patient-derived glioma models included SU-DIPG-13fl, SU-DIPG-17, SU-DIPG-19, and SU-DIPG-92. Patient characteristics for these models are detailed in Supplemental Information 1. Throughout the culture period, all cultures were subjected to monitoring for authenticity via short tandem repeat (STR) fingerprinting and routine mycoplasma testing was conducted. The glioma cultures were cultivated as neurospheres in serum-free medium composed of DMEM (Invitrogen), Neurobasal(-A) (Invitrogen), B27(-A) (Invitrogen), heparin (2 ng ml−1), human-bFGF (20 ng ml−1) (Shenandoah Biotech), human-bEGF (20 ng ml−1) (Shenandoah Biotech), human-PDGF-AA (10 ng ml−1) (Shenandoah Biotech), and human-PDGF-BB (10 ng ml−1) (Shenandoah Biotech). For *in vitro* experiments, the neurospheres were dissociated using TrypLE (Gibco) for seeding.

### Animal Models

Homozygous ChAT-IRES-Cre mice (created by Dr. Bradford Lowell and obtained from The Jackson Laboratory, strain 006410) were bred with homozygous Ai230 mice (Drinnenberg et al., in preparation) Optogenetic experiments were performed on animals with the genotype ChAT-IRES-Cre^+/wt^ x Ai230^flx/wt^, which enabled optogenetic control selectively for cholinergic neurons in immunocompetent mice. All mice used were genotyped at postnatal day 10. For tumour studies, glioma cell implantation was conducted using an electroporated, engineered H3.3K27-altered MADR^36^ model. To replicate results obtained from this immunocompetent tumour model in patient-derived glioma cells, viral vectors (specified below) were injected into 4-week-old NSG mice (NOD-SCID-IL2R-gamma chain-deficient, The Jackson Laboratory), allowing optogenetic control of the laterodorsal tegmentum nucleus and pedunculopontine nucleus in immunodeficient mice, followed by xenografting of patient-derived DIPG cells (SU-DIPG-17) after 3 weeks of virus expression. All animal experiments were conducted in accordance with protocols approved by the Stanford University Institutional Animal Care and Use Committee (IACUC) and performed in accordance with institutional guidelines. Animals were housed according to standard guidelines with unlimited access to water and food, under a 12 h light: 12 h dark cycle, a temperature of 21 °C and 60% humidity. For brain tumour allograft or xenograft experiments, the IACUC has a limit on indications of morbidity (as opposed to tumour volume). Under no circumstances did any of the experiments exceed the limits indicated and mice were immediately euthanized if they exhibited signs of neurological morbidity or if they lost 15% or more of their initial body weight.

### Generation of Transgenic Ai230 Mice

Ai230 (full strain name TIGREtm230(TIT2L-XCaMPG-WPRE-ICL-ChRmine-TS-oScarlet-Kv2.1-ER-IRES2-tTA2-WPRE_hyg)Hze) (Drinnenberg et al., in preparation) is a new TIGRE2.0 transgenic reporter^69^, which provides Cre-dependent expression of soma-targeted ChRmine-oScarlet and Cre- and tTA-dependent expression of XCaMPG. Ai230 mice contain a modified TIGRE genomic locus that contains: (5’ to 3’): A Frt3 site, two tandem copies of the chicken beta-globin HS4 insulator element, a Tet responsive 2 promoter comprised of seven repeats of TRE binding sites and a minimal CMV promoter (based on that in Clontech’s pTRE2-hyg vector), a loxP site, a stop cassette (with stops in all three frames linked to a synthetic pA-hGH pA-PGK pA unit), a loxP site, the coding sequence for XCaMP-G, a woodchuck post-transcriptional regulatory element (WPRE), a bGH pA, two tandem copies of the chicken beta-globin HS4 insulator element, a CAG promoter which consists of the CMV enhancer fused to the chicken beta-actin promoter, a lox2272 site, a stop cassette (with stops in all three frames linked to a synthetic pA-hGH pA-TK pA unit), a lox2272 site, the coding sequence of ChRmine, a membrane trafficking signal (TS), oScarlet, Kv2.1, a woodchuck post-transcriptional regulatory element (WPRE), a bGH pA, an IRES2 sequence, tetracycline-transactivator 2 (tTA2), a woodchuck post-transcriptional regulatory element (WPRE), a bGH pA, a PGK promoter, one domain of the hygromycin resistance gene, a mRNA splice donor sequence, a Frt5 site. The locus was generated by recombinase mediated cassette exchange into a previously made docking site integrated into the TIGRE locus.

### Generation of H3.3 K27M MADR cells

The H3.3 K27M MADR tumor cell cultures were generated using the same technique as previously described^36,73^. Briefly, Gt(ROSA)26Sortm4(ACTB-tdTomato,-EGFP)Luo/J and Gt(ROSA)26Sortm1.1(CAG-EGFP)Fsh/Mmjax mice (JAX Mice) were crossed with wild-type CD1 mice (Charles River) to produce heterozygous mice. Male and female embryos between E12.5 and E15.5 were subjected to *in utero* electroporations. Pregnant dams were individually housed, and pups remained with their mothers until P21 in the institutional animal facility (Stanford University). The MADR tumor cell line used here was generated by dissociating and sorting GFP+ tumor cells from female heterozygous mTmG mice. Subsequently, MADR cultures were maintained as neurospheres in serum-free medium composed of DMEM (Invitrogen), Neurobasal(-A) (Invitrogen), B27(-A) (Invitrogen), heparin (2 ng ml−1), human bFGF (20 ng ml−1) (Shenandoah Biotech), human bEGF (20 ng ml−1) (Shenandoah Biotech), human PDGF-AA (10 ng ml−1) (Shenandoah Biotech), insulin (Sigma-Aldrich), and 2-mercaptoethanol (Sigma-Aldrich).

### Stereotaxic Surgery, Ferrule Placement, and Viral Vectors

Mice were anesthetized with 1-4% isoflurane and placed in a stereotaxic apparatus. For all optogenetic experiments, mice were unilaterally implanted with optical fibers above either the laterodorsal tegmentum nucleus (LDT) or the pedunculopontine nucleus (PPN) on the right side. Optical fibers were secured with stainless steel screws (thread size 00-90 x 1/16, Antrin Miniature Specialties), C&B Metabond, and light-cured dental adhesive cement (Geristore A&B paste, DenMat). For optogenetic stimulation of NSG mice, 1 µl of AAV-ChR2::eYFP or AAV-eYFP viral vectors were unilaterally injected using Hamilton Neurosyringe and Stoelting stereotaxic injector over 5 minutes. Coordinates for viral injections and optic ferrule placements were measured from bregma. Viral vectors were injected, and an optic ferrule was placed into laterodorsal tegmentum nucleus (LDT) at coordinates, AP=-5.0mm, ML=+0.5mm, DV=-3.15mm, and for pedunculopontine nucleus (PPN) stimulations at AP= −4.70mm, ML=+1.25mm, DV=-3.50mm.

### Allografting and Xenografting

Male and female mice were used in cohorts equally. For optogenetic stimulation studies, MADR cultures (*H3.3 K27 MADR line 1, n = 200.000 cells per mice*) or patient-derived DIPG cultures (*SU-DIPG-17,* n = 400.000 cells per mice) were injected into the thalamus or pontine region. A single-cell suspension of all cultures was prepared in sterile culture medium immediately before surgery. Animals at P28–P35 were anaesthetized with 1–4% isoflurane and placed on stereotactic apparatus. Under sterile conditions, the cranium was exposed via a midline incision and a 31-gauge burr hole made at exact coordinates. For thalamus injections the coordinates were as follows: AP=-1.0mm (from bregma), ML=+0.8mm, DV=-3.5mm. For pontine injections coordinates were AP=-0.8 (from lambda), ML=−1.0mm, DV=−5.0mm. Cells were injected using a 31-gauge Hamilton syringe at an infusion rate of 0.5 μl min−1 with a digital pump. At completion of infusion, the syringe needle was allowed to remain in place for a minimum of 5 minutes, then manually withdrawn. The wound was closed using 3M Vetbond (Thermo Fisher Scientific) and treated with Neo-Predef with Tetracaine Powder.

### Viral Vectors

For the optogenetic stimulation of cholinergic midbrain nuclei (LDT and PPN) in immunodeficient NSG mice, expression of channelrhodopsin in each nucleus was achieved by injection of AAV-hSyn-hChR2(H134R)::eYFP (virus titer= 1.4×10^12^ gc/ml), and eYFP control by injection of AAV-hSyn::eYFP (virus titer= 1.5×10^12^ gc/ml). Both viral vectors were obtained from Stanford University Gene Vector and Virus Core.

### Optogenetic Stimulation

Optogenetic stimulations were performed 1 week after optic ferrule implantation. For allograft experiments in ChAT-IRES-Cre^+/wt^ x Ai230^flx/wt^ mice, freely moving animals were connected to a 595 nm high-power LED system with a monofiber patch cord achieving stimulation of ChRmine. For xenograft experiments in immunodeficient NSG mice, a 473 nm diode-pumped solid-state laser system was used to achieve stimulation of ChR2. Cholinergic neuron stimulation, for both neuronal cell bodies and axon terminals, was performed at 20 Hz, ten 15 ms pulses of 595 nm light delivery every 5 seconds at a light power output of 10mW from the tip of the optic fiber (200 µm core diameter, NA=0.22 - Doric lenses). Optogenetic stimulation session lasted for 30 minutes. Animals were injected intraperitoneally with 40 mg/kg EdU (5-ethynyl-2’-deoxyuridine; Invitrogen, E10187) before the session, and perfused 3 hours (for OPC response analysis) or 24 hours (for glioma cell proliferation analysis) after the start of the stimulation. The effectiveness of optogenetic stimulation in the ChAT^+/wt^ x Ai230^flx/wt^ model was validated through cFos staining, as depicted in Supplementary Figure 1. In Ai230 models, light delivery results in stimulation. Thus, we designated the group with the "laser on" as the stimulated cohort, while the non-stimulated group did not receive light ("laser off").

### Pharmaceutical Antagonism of Muscarinic Receptors M1 and M3

To assess a potential effect of muscarinic receptors M1 (CHRM1) and M3 (CHRM3) on the proliferative effect of cholinergic neuronal activity in vivo, ChAT-IRES-Cre^+/wt^ x Ai230^flx/wt^ mice were allografted with the MADR model as above and blind randomized to a treatment group. Four weeks post-allograft, mice were treated with intraperitoneal administration of the M1 blocker VU 0255035 (10 mg kg−1; Tocris) or 4-DAMP (10 mg kg−1; Tocris) and controls treated with equivalent volume of vehicle. Administration of with the drug or vehicle was done 30 minutes before the beginning of the optogenetic stimulation session of each mice. For immunohistological analysis of glioma cell proliferation, mice were perfused 24 hours after optogenetic stimulation.

### Cerebral Slice Conditioned Media of LDT and PPN

ChAT-IRES-Cre^+/wt^ x Ai230^flx/wt^ mice aged 4 weeks (P28 to P30) were utilized to collect conditioned media from activated cholinergic neurons in either the LDT or PPN. Brief exposure to isoflurane induced unconsciousness in the mice before immediate decapitation. Extracted brains (cerebrum) were inverted and placed in an oxygenated sucrose cutting solution, then sliced at 300μm to target the region of the LDT or PPN. Slices (n=4 per mouse) were transferred to ACSF and allowed to recover for 30 minutes at 37°C, followed by an additional 30 minutes at room temperature. After recovery, the slices were transferred to fresh ACSF and positioned under a red-light LED using a microscope objective. The optogenetic stimulation paradigm consisted of 20-Hz pulses of red light for 30 seconds on, followed by 90 seconds off, repeated over a period of 30 minutes. Conditioned medium from the surrounding area was collected and stored frozen at −80°C. Stimulated slices were postfixed in 4% paraformaldehyde (PFA) for 30 minutes before cryoprotection in 30% sucrose solution for 48 hours. Successful stimulation of cholinergic neurons in each area was validated through cFos staining.

### EdU Incorporation Assay

EdU staining of glioma monocultures or glioma-neuron co-cultures was performed on glass coverslips in 96-well plates which were pre-coated with poly-l-lysine (Sigma) and laminin (Thermo Fisher Scientific). Neurosphere cultures were dissociated with TrypLE and plated onto coated coverslips with growth factor-depleted medium. Acetylcholine (0.5μM to 5μM, Tocris), VU 0255035 (10μM, Tocris), 4-DAMP (10μM, Tocris) and vehicle (DMSO) were added for specified times with 4 μM EdU. After 24 h the cells were fixed with 4% PFA in PBS for 20 min and then stained using the Click-iT EdU kit and protocol (Invitrogen) and mounted using Prolong Gold mounting medium (Life Technologies). Confocal images were acquired on a Zeiss LSM980 using Zen 2011 v8.1. Proliferation index was determined by quantifying the fraction of EdU-labelled cells divided by DAPI-labelled cells (monoculture experiments), or HNA-labeled cells (co-culture experiments) using confocal microscopy at 20× magnification. Quantification of images was performed by a blinded investigator.

### Migration Assay

3D migration experiments were performed as previously introduced^74^ with some modifications. Briefly, 96-well flat-bottomed plates (Falcon) were coated with 2.5μg per 50μl laminin per well (Thermo Fisher) in sterile water. After coating, a total of 200μl of culture medium per well was added to each well. A total of 100μl of medium was taken from 96-well round bottom ULA plates containing ∼200μm diameter neurospheres, and the remaining medium including neurospheres was transferred into the pre-coated plates. Images were then acquired using an Evos M5000 microscope (Thermo Fisher Scientific) at time zero, 24, 48, and 72 hours after encapsulation. Image analysis was performed using ImageJ by measuring the diameter of the invasive area. The extent of cell migration on the laminin was measured for six replicate wells normalized to the diameter of each spheroid at time zero and the data is presented as a mean ratio for three biological replicates.

### Neuron-Glioma Co-Culture

For EdU incorporation assays, neurons were isolated from CD1 mice (The Jackson Laboratory) at P0 using he Neural Tissue Dissociation Kit Postnatal Neurons (Miltenyi), and followed by the Neuron Isolation Kit, Mouse (Miltenyi) per manufacturer’s instructions. After isolation, 200.000 neurons were plated onto circular glass coverslips (Electron Microscopy Services) pre-coated with poly-l-lysine (Sigma) and mouse laminin (Thermo Fisher Scientific). Neurons were cultured in BrainPhys neuronal medium containing B27 (Invitrogen), BDNF (10 ng ml−1, Shenandoah Biotech), GDNF (5 ng ml−1, Shenandoah Biotech), TRO19622 (5 μM; Tocris) and β-mercaptoethanol (Gibco). The medium was replenished on days *in vitro* (DIV) 1 and 3. On DIV 5, fresh medium was added containing 70.000 glioma cells and incubated for 48 h. After 48h incubation, EdU (10 μM) with or without the acetylcholine (0.5-5 μM, Tocris) was added and incubated for a further 24h. Following incubation, the cultures were fixed with 4% paraformaldehyde (PFA) for 20 min at room temperature and stained for immunofluorescence analysis. For EdU analysis, cells were stained using the Click-iT EdU Cell Proliferation kit (Thermo Fisher Scientific, C10337), before staining with primary antibodies mouse anti-human nuclei clone 235-1 (1:250; Millipore, MAB1281) and rabbit anti-microtubule-associated protein 2 (MAP2; 1:500, EMD Millipore, AB5622), overnight at 4 °C. Following washing, slips were incubated in secondary antibodies, Alexa 488 donkey anti-mouse IgG (1:500, Jackson Immuno Research) and Alexa Fluor 555 donkey anti-rabbit (1:500, Invitrogen) and mounted using ProLong Gold Mounting medium (Life Technologies). Images were collected on a Zeiss LSM980, and proliferation index determined by quantifying percentage EdU-labelled glioma cells over total glioma cells (HNA immunopositivity to identify glioma cells).

### Human iPSC-Derived Cholinergic Neurons

Human induced pluripotent stem cells (iPSCs) of a 12-year-old male healthy donor (CW20032, Elixirgen) maintained under feeder-free conditions in a 96-well plate (20.000 cells per well) were rapidly differentiated to a cholinergic phenotype with Sendai virus mediated delivery of synthetic neurogenic transcription factor (CH-SeV, Elixirgen) as previously described^75^. After successful morphological differentiation into cholinergic neurons at day 7, fresh medium (cholinergic neuron maintenance media, CH-MM, Elixirgen) was added containing 20.000 glioma cells (“SU-DIPG-17”) and incubated for 48 h. After 48h incubation, EdU (4 μM) with or without VU 0255035 (10μM, Tocris) or 4-DAMP (10μM, Tocris) was added and incubated for a further 24h. Following incubation, the cultures were fixed with 4% paraformaldehyde (PFA) for 20 min at room temperature and further staining was processed as above described.

Primary antibodies mouse anti-nestin (1:500; Abcam, ab6320), rabbit anti-choline acetyltransferase (ChAT; 1:500, Abcam, ab181023) and goat anti-microtubule-associated protein 2 (MAP2; 1:500, Abcam, ab302488) were used. For synaptic puncta staining, fresh medium was added containing 50.000 glioma cells (“SU-DIPG-13fl”) expressing PSD95–RFP (as generated in Taylor et al.^8^) and incubated for 72h. After fixation with 4% PFA, cholinergic neuron-to-glioma co-culture coverslips were incubated in blocking solution (3% normal donkey serum, 0.3% Triton X-100 in TBS) at room temperature for 1h. Primary antibodies guinea pig anti-synapsin1/2 (1:500; Synaptic Systems, 106-004), rabbit anti-RFP (1:500; Rockland, 600-401-379), mouse anti-nestin (1:500; Abcam, ab6320), and goat anti-MAP2 (1:500, Abcam, ab302488) diluted in diluent (1% normal donkey serum in 0.3% Triton X-100 in TBS) and incubated at 4 °C overnight. Following washing, the slides were incubated in secondary antibody (Alexa 555 donkey anti-rabbit IgG, Invitrogen; Alexa 405 donkey anti-guinea pig IgG; Alexa 647 donkey anti-mouse IgG and Alexa 594 donkey anti-goat IgG all used at 1:500, Jackson Immuno Research) overnight at 4 °C. Following washing, coverslips were mounted using ProLong Gold Mounting medium (Life Technologies). Images were collected on a Zeiss LSM980 confocal microscope using a 63× oil-immersion objective. Co-localization of synaptic puncta images were performed using a custom ImageJ (v.2.1.0/153c) processing script. In brief, the quantification determines co-localization of presynaptic synapsin and postsynaptic PDS95–RFP within a defined proximity of 1.5 μm. Background fluorescence is removed using rolling ball background subtraction and peaks detected using imglib2 DogDetection plugin which determines the region of interest for each channel. The percentage of total glioma ROIs that are within 1.5 μm of a neuron ROI is reported. The script was implemented in ImageJ (v.2.1.0/153c).

### Immunohistochemistry

All mice were anesthetized with intraperitoneal injections of 2.5% Avertin (tribromoethanol; Sigma-Aldrich, T48402), and transcardially perfused with 20 ml 0.1M phosphate buffer saline (PBS). Brains were postfixed in 4% paraformaldehyde (PFA) overnight at 4°C before cryoprotection in 30% sucrose solution for 48 hours. For sectioning, brains were embedded in optimum cutting temperature (OCT; Tissue-Tek) and sectioned coronally at 40 µm using a sliding microtome (Leica, HM450). For immunohistochemistry, brain sections were stained using the Click-iT EdU cell proliferation kit (Invitrogen, C10339 or C10337) according to manufacturer’s protocol. Tissue sections were then stained with antibodies following an incubation in blocking solution (3% normal donkey serum, 0.3% Triton X-100 in tris buffer saline) at room temperature for 30 minutes. Mouse anti-human nuclei clone 235-1 (1:100; Millipore, MAB1281), rabbit anti-ChAT (1:500; Abcam, ab223346), goat anti-Pdgfra (1:500; R&D Systems, AF1062), rat anti-MBP (1:200; Abcam, ab7349), chicken anti-mCherry (1:1000; Abcam, ab205402), or rabbit anti-cfos (1:500; Santa Cruz Biotechnology, sc-52) were diluted in 1% blocking solution (1% normal donkey serum in 0.3% Triton X-100 in TBS) and incubated overnight at 4°C. All antibodies have been validated in the literature for use in mouse immunohistochemistry. The following day, brain sections were rinsed three times in 1x TBS and incubated in secondary antibody solution for 2 hours at room temperature. All secondary antibodies were used at 1:500 concentration including Alexa 488 anti-rabbit, Alexa 488 anti-mouse, Alexa 488 anti-chicken, Alexa 594 anti-chicken, Alexa 647 anti-goat, Alexa 647 anti-rat, Alexa 647 anti-rabbit, Alexa 405 anti-guinea pig (all Jackson ImmunoResearch), and Alexa 555 anti-rabbit (Invitrogen). Sections were then rinsed three times in 1x TBS and mounted with ProLong Gold (Life Technologies, P36930).

### Confocal Microscopy and Quantification

All image analysis were performed by experimenters blinded to the experimental conditions or genotype. Cell quantification within allografted or xenografted tumors was conducted by acquiring z-stacks using a Zeiss LSM980 scanning confocal microscope (Carl Zeiss). A 1-in-6 series of coronal brain sections were selected, with 4 consecutive slices (40µm thickness) analyzed in the grafted brain area (thalamus, or pons). Brain tissue damaged during perfusion or tissue processing was excluded from histological analysis. Tumor cells were identified as GFP^+^ (MADR allografts) or HNA^+^ (patient-derived xenografts) and quantified in each field to determine the proliferation index, calculated as the percentage of GFP^+^ cells co-labeled with EdU. OPCs were identified by PDGFRa staining and quantified as the percentage of PDGFR^+^ cell co-labeled with EdU.

### Western Blotting

For western blot analysis of NLGN3 in conditioned media, samples were prepared by adding LDS buffer and β-mercaptoethanol, heated, and loaded onto Tris-acetate gels. After electrophoresis and transfer onto PVDF membranes, the membranes were blocked and incubated with primary antibodies against the NLGN3 ectodomain (Abcam #ab192880). Following incubation with secondary antibody and washing, the membranes were developed using chemiluminescent substrate (Thermo Scientific #34580).

### Measurement of BDNF and acetycholine in Conditioned Media

The concentration of BDNF (R&D Systems #DBNT00) and acetylcholine (Sigma-Aldrich # MAK056) in conditioned media was measured using ELISA. We compared the concentrations of conditioned media obtained from non-stimulated, LDT-stimulated, and PPN-stimulated brain slices as described above. Protocols were performed according to the manufacturer’s instructions. The optical density was measured at 450 nm with an absorbance microplate reader (Molecular Devices, SpectraMax M3). The total concentrations were determined as pg/mL from the standard curve. To confirm the specificity of the respective kit, medium and lysis buffer without protein extract were used as negative controls.

### Single-Cell RNA Sequencing Analysis of Published Data Cell filtering

We excluded cells with a low number of detected genes, using a cutoff of 2000 genes for smart-seq2 data and a cutoff of 1000 genes for the other types of sequencing data.

### Gene filtering

In analyses that necessitated gene filtering, we kept the 7000 genes with the highest mean expression across cells.

### Normalization

UMI counts were converted to counts per million (CPM). Each entry in the expression matrix was then normalized according to 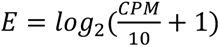. The same normalization was used for transcripts per million (TPM) values. The values were divided by 10 as the actual complexity is assumed to be in the realm of about 100,000 and not 1 million as implied by the CPM and TPM measures.

### Centering

To illustrate intra-tumor heterogeneity in gene expression (Supplementary Figure 4A), centering was performed individually for each sample. This involved subtracting each gene’s mean expression value across all cells within the sample. Cells within each sample were then sorted based on their scores for the cholinergic genes using the sigScores function available at https://github.com/jlaffy/scalop.

For the pseudo-bulk data analysis (Figure 4A), expression values were first averaged across malignant cells within each sample. Subsequently, log-normalization was applied, but no centering was performed.

### Correlation with cell states

Utilizing the OPC-like signature outlined by Neftel et al.^44^, we conducted the following analyses for both H3K27M DGM samples and medulloblastoma samples:

i. We computed the correlations within each sample between the cells’ OPC-score and the expression values of each selected cholinergic gene.
ii. We calculated the correlations across samples within each study, between the pseudo-bulk sample OPC scores and the pseudo-bulk gene expression levels.

These correlation values were then averaged across samples and studies, respectively (see Figure 4B and Supplementary Figure 4B).

Additionally, we assessed correlations across samples in other tumor types using more cell-state signatures (Neftel et al., 2019^44^; Gavish et al., 2023^47^) (Figure 4C). Significant correlations (P < 0.05 after FDR correction) were considered.

### Lineage vs stemness analysis

Focusing on the H3K27M DGM data published in the Filbin et al. 2018^13^ study, we computed a ‘stemness score’ for each cell. This score is determined by subtracting either the cell’s OC score or its AC score from its OPC score, whichever is higher (the maximum of the two). We also calculated a ‘lineage score’ for each cell, defined as the maximum between the OC and AC scores. In cases where the AC score is higher, it is multiplied by −1. If both AC and OC scores are negative, a 0 value is assigned with some jitter. Additionally, we identified cells with a centered CHRM1 value > 1 and cells with a centered CHRM3 value > 1. All scoring and centering procedures were performed per sample.

### Statistical Analysis

Gaussian distribution was confirmed using the Shapiro-Wilk test. Parametric data were analyzed with unpaired two-tailed Student’s t-tests or one-way ANOVA with Tukey’s post hoc tests. Significance was set at p < 0.05. The used statistical test is indicated in the figure legend. GraphPad Prism 10 was used for statistical analyses and data illustrations.

## ACKNOWLEDGMENTS

We express our heartfelt gratitude to all the children and their families who generously donated tumor tissue for research. We want to emphasize that this research would not have been feasible without their invaluable donations. Additionally, we extend our thanks to all members of the Monje Lab for fruitful discussions and technical assistance. Funding: National Institute of Neurological Disorders and Stroke (R01NS092597 to MM), NIH Director’s Pioneer Award (DP1NS111132 to MM), National Cancer Institute (P50CA165962, R01CA258384, U19CA264504 to MM), Cancer Research UK (to MM), Cancer Grand Challenges (OT2CA278688, CGCATF-2021/100012), Gatsby Charitable Foundation (Gatsby Initiative in Brain Development and Psychiatry, to M.M. and K.D.), Oscar’s Kids Foundation (to M.M.), McKenna Claire Foundation (to M.M.), Will Irwin Research Fund of the Pediatric Cancer Research Foundation (to M.M.), Yuvaan Tiwari Foundation (to M.M.), Austin Strong Foundation (M.M.), Avery Huffman DIPG Foundation (to M.M.), Chadtough Defeat DIPG (to M.M.),Maternal and Child Health Research Institute at Stanford University Postdoctoral Award (to BY), Dean’s Postdoctoral Fellowship at Stanford University (to BY), Damon Runyon Cancer Research (to K.R.T.), Stanford Maternal & Child Health Research Institute (to K.R.T.), DoD grant HT9425-23-1-0269 (to J.B.), NIH grant R33CA236687 (to J.B.), American Cancer Society grant RSG-16-217-01-TBG (to J.B.).

## AUTHOR CONTRIBUTIONS

Conceptualization, R.D. and M.M.; Methodology, R.D., A.D., B.Y., K.S., K.R.T., A.E.A., D.R.P., J.B., C.R., L.S., T.L.D., B.T., and H.Z.; Software, A.G.; Formal Analysis, R.D., A.G., A.R., and Y.S.K.; Investigation, R.D., A.R., R.M., Y.S.K., P.J.W., A.R., and E.T.; Resources, A.D., K.S., K.R.T., J.B., K.D., and M.M.; Writing – Original Draft, R.D. and M.M.; Writing – Review & Editing, all authors; Visualization, R.D. and A.G.; Supervision, M.M.; Funding Acquisition, M.M.

## RESOURCE AVAILABILITY

### Lead contact

Further information and requests for resources and reagents should be directed to and will be fulfilled by the lead contact, Michelle Monje.

### Materials availability

This study did not generate new unique reagents. The availability of the Ai230 mouse line will be instructed in Drinnenberg et al. (in preparation) and the generation of the MADR tumor line is detailed described in Kim et al. (Cell, 2019).

### Data and code availability

All analyzed single-cell and single-nucleus RNA sequencing data have been deposited by the study groups that generated the datasets. Any additional information required to reanalyze the data reported in this paper is available from the lead contact upon request.

